# NetTracer3D Enables User-Friendly Analysis of Diverse Microscopic and Medical 3D Datasets

**DOI:** 10.64898/2026.03.25.714104

**Authors:** Liam Mclaughlin, Mia Curic, Siddhartha Sharma, Jorge Villazon, Rebecca J. Salamon, Manako Yamaguch, Maria Luisa S. Sequeira-Lopez, Philippa R. Kennedy, Riley C. Lyons, Lingyan Shi, R. Ariel Gomez, Sanjay Jain

**Affiliations:** Washington University School of Medicine, Department of Medicine, St. Louis, MO; Shu Chien-Gene Lay Department of Bioengineering, Jacob School of Engineering, UC San Diego, CA 92093; Department of Pediatrics, School of Medicine, University of California, San Diego, La Jolla, CA 92093, USA; Department of Pediatrics, Child Health Research Center, University of Virginia School of Medicine; Department of Medicine, University of Minnesota Medical School, Minneapolis, MN, USA; Washington University School of Medicine, Department of Pathology & Immunology, St. Louis, MO; Washington University School of Medicine, Department of Pediatrics, St. Louis, MO

## Abstract

Recent advances in high-resolution imaging and spatial transcriptomics have enabled reconstruction of complex 3D tissue maps, providing unprecedented insights into cellular connectivity, organization, and tissue architecture. However, standardization challenges hinder integration, sharing, and analysis of these datasets across research communities. We developed NetTracer3D to simplify three-dimensional image analysis across diverse datasets. NetTracer3D is an integrated tool for defining, processing, and sharing 3D tissue maps with standardized data formats and interactive exploration capabilities. It provides three broadly applicable network analysis modalities: Connectivity networks for analyzing functional tissue units or cells connected via secondary structures such as nerves or vasculature; Branch Adjacency and Branchpoint networks for converting branched anatomical structures into analyzable representations; and Proximity networks for grouping structures by spatial relationships to identify cellular organization patterns. We demonstrate several use cases applying NetTracer3D to analyze multidimensional data from CODEX and label free Raman spectroscopy, multiscalar data encompassing subcellular and anatomical scales and a range of modalities. NetTracer3D was able to characterize neural relationships between functional tissue units in human and mouse kidneys and mouse bronchi. Branchpoint networks were used to identify vascular defects in human brain angiogram and define the innervation structure of a lymph node. Finally, we demonstrate how proximity networks characterize the tumor microenvironment in 3D light sheet cancer images and auto-detect cellular neighborhoods in multiplexed 2D CODEX datasets. Beyond network creation, NetTracer3D enables analysis, spatial statistics, and visual analytics tailored for volumetric tissue data. By establishing interoperable formats and analysis workflows, this work provides accessible and reproducible analytical tools for 3D spatial biology, enabling new discoveries of relationships between structure and physiology.

## Main

Analysis of network structures has played a key role in explaining many biological interactions, including protein-protein, signaling, regulation, integration of physiological systems and metabolic pathways^1^. While such methods are common for explaining interactions on biochemical levels such as involving genes and proteins^2^, NetTracer3D represents an early foray in their application in 3D imaging. Current methods of cell clustering often rely on arbitrarily sized groups of nearest neighbors, which can fail to include, or otherwise over-include, objects in cellular neighborhoods^3^. Furthermore, while such use of automated clustering methods is more common at cellular scales, there remains a need to characterize connectivity amongst niche environments of functional tissue units (FTU)^4^ in order to better describe biological systems across different scales of organization, which has limited our understanding of niche biology^5^. Three-dimensional (3D) spatial data provides unique insights into how tissues are organized and form connectivity networks that regulate homeostasis and disease. While new open-source tools have been developed for 3D analysis^6^, several factors including accessibility, high cost, and complex workflows have limited their adoption^7^. Furthermore, many of the current open-source tools lack a graphical user interface (GUI), requiring coding knowledge and expertise to install and use properly.

NetTracer3D provides three broadly applicable network analysis modalities for diverse imaging applications. First, **Connectivity networks** enable analysis of functional tissue units or cells connected via secondary structures such as nerves or vasculature. In this paper, we first explore how connectivity networks can be used to characterize neurovascular connectivity kidney FTUs and lung bronchi, however this applicability extends to virtually any segmented object. Network analytics of the structures and communities that are formed offer a simplistic way to perform quantitative comparative analysis between separate 3D datasets, offering new avenues to characterize how pathological states affect tissues.

Second, **Branch Adjacency and Branchpoint networks** convert branched anatomical structures, including nervous, vascular, and lymphatic systems, into analyzable network representations. Where Connectivity Networks require the presence objects to be connected, Branch Networks are broadly applicable, rapidly deconstructing a three dimensional nervous or vascular structure into a simplistic network dataset that can be immediately and broadly quantified across a plethora of network statistics. This allows experimental conclusions to be drawn solely from single-channel images of nerves or blood vessels.

Third, **Proximity networks** group structures by spatial nearness, particularly useful for analyzing cellular organization. Although similar neighborhood grouping approaches have been applied to cellular data^8^, NetTracer3D is unique in its ability to characterize such structures in 3D space, a critical development in the wake of the development of novel 3D multiplexing techniques^9^.

By developing NetTracer3D, we have provided a user-friendly, open access platform designed to enhance utility and applicability of large-scale volumetric analysis. NetTracer3D is compatible across a variety of image files and data types, with features not easily available from existing commercial vendors such as Imaris and Amira. NetTracer3D empowers users as it is an open-source 2D/3D Network Analysis platform for microscopy and medical images, featuring a user-friendly GUI that enables end-to-end image segmentation, processing, and analysis. It supports end-to-end analysis pipelines, beginning with segmentation, continuing to network generation, and finally providing rapid and broadly applicable ways to quantify networks to reach experimental conclusions for 3D datasets.

## Results

### Overall workflow of NetTracer3D

To demonstrate capabilities of NetTracer3D for 3D image analysis, we first provide an overview of its basic functions by using a real-world example of feeding behavior of slime mold and infrastructure networks and then various applications to 2D and 3D data across macro-and-microscopic modalities in biological systems. The overall workflow for NetTracer3D works as follows. After the acquisition and necessary preprocessing of a microscopic/radiographic image, the image is segmented for node objects (to be grouped into a network)^10^, either by creating a binary mask for structures of interest (which may be performed within NetTracer3D, or another segmenting software), or creating a mask of individually labeled objects, such as labeling cells with a software such as Cellpose3^11^. NetTracer3D next offers a plethora of tools for refining segmentation masks, including assigning labels to discrete objects via watershedding^12^ and for morphological analysis of said labeled objects, such as directly analyzing clustering arrangements with a sophisticated Ripley’s analysis algorithm^13^. Node objects may further be grouped into networks through three primary modalities: The first is the ‘connectivity’ network, where objects are characterized by their shared connectivity through some other medium, such as evaluating shared nerves or vasculature. The second are branch networks, where branched objects such as nerves or vessels are directly discretized into their branched subcomponents, and grouped into a network of adjacent branches or branchpoints. The third modality is the proximity network, where objects are connected to nearby objects, which is ideal for cellular datasets. Generated networks may then be directly grouped into communities and quantified with networkX^14^.

Figure 1 demonstrates an example of an end-to-end analysis using NetTracer3D, starting with a raw image and ending with network statistics using an example from slime mold and similarity between plasmodium arrangements and real-world railway networks in Tokyo^15^ and extracting network connectivity data. **Step 1:** Identify objects to be used as nodes (to be grouped into a network, in this case, the large feeding structures in the slime mold) and optionally, as second to use as edges (connections in a network, in this case, the thin psuedopods of the slime mold)^10^. **Step 2:** The image must be imported into the nodes channel for segmentation prior to network generation (Figure 1a). Objects of interest can be segmented prior in other software, or through NetTracer3D’s own segmentation GUI (Figure 1b) which implements a machine learning random-forest classifier in a similar manner as the ImageJ labkit extension^16^. **Step 3:** Import desired nodes and edges into their respective channels in the GUI. **Step 4:** After thresholding out noise (optional), call the ‘Calculate Connectivity Network’ from the ‘Calculate Network’ submenu to convert the segmented channels into network data (Figure 1c). Once the network has been generated, the interactive GUI allows for direct exploration and evaluation of specific objects. 3D image viewing of the images and any generated overlays is then available through bundling with napari^17^. **Step 5:** The generated networks undergo comprehensive analysis using NetworkX^18^, a Python-based package for network analysis that has been extensively validated in biomedical applications including microscopy image analysis and vascular network analysis. The GUI features both an interactive image viewer that is linked to an interactive network graph view, allowing easy discovery and visual characterization of nodes of interest. Statistical characterization includes community clustering (Figure 1d) and computation of fundamental graph-theoretic measures that describe network topology both at the nodal level (e.g., clustering coefficients, closeness centrality, eigenvector centrality, betweenness centrality, degree centrality) and as network-wide distributions (Figures 1e**, f**). These metrics provide quantitative insights into network structure and connectivity patterns that are particularly relevant for understanding cellular organization in microscopic images^3^. In addition to these calculations, any statistical table generated in NetTracer3D that describes the nodes with a quantifiable parameter may then be used as a threshold to isolate specific nodes in the image for further visualization or analysis (Figure 1e).

**Figure 1.**
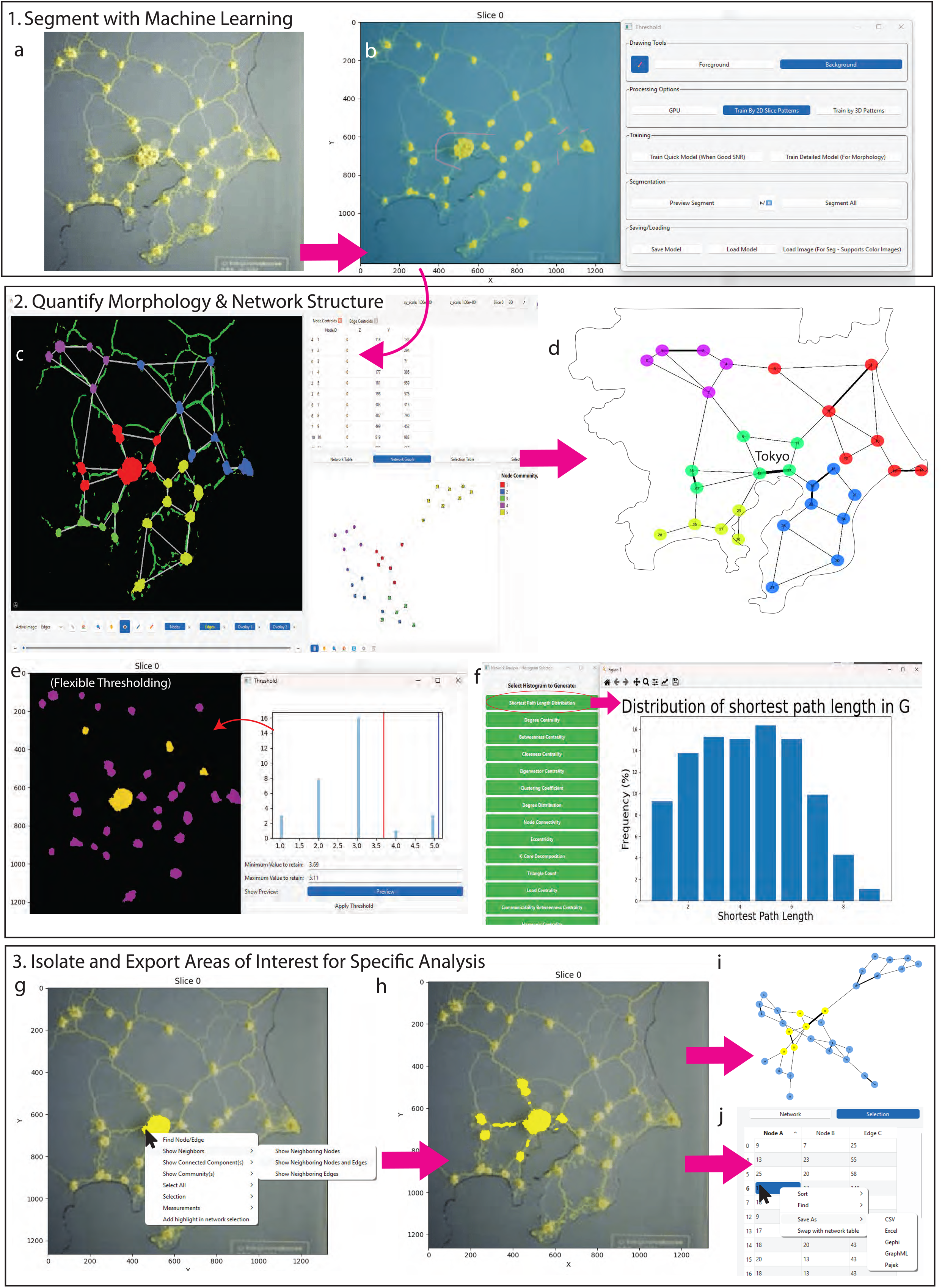
NetTracer3D Example Pipeline – The Slime Mold. **a**. The slime mold from Tero et al^15^ that had been grown to resemble the Tokyo railway map. **b.** Example of NetTracer3D’s segmentation window. Foreground and background regions are denoted by the user with green or red scribbles, respectively. Earlier versions used this data to train a random forest classifier model^67^, although this was updated to use Lightgbm instead as of version 1.5.5. In this instance, the ‘node’ objects are being segmented, predicted as foreground with the yellow overlay, while the predicted background is shown blue. **c.** NetTracer3D’s user interface. The central image viewer window shows the segmented nodes (magenta) and edges (green), with optional background overlays. Loading, analysis, and processing can be executed from the upper left set of menus, while the lower left menu bar allows interaction with the image in various ways. In the upper-right is a window that will be populated by any generated data tables; in this instance, showing centroids of node objects. In the lower right is the network menu, which shows pairs of nodes and the edge-ids that connect them (if any). After segmentation of the nodes and desired edge objects (for ‘connectivity networks’, in this case), NetTracer3D assigns unique labels to nodes and uses them to search for nearby edges to yield network pairs. **d**. A network graph with Louvain communities generated by NetTracer3D, manually overlayed on a tracing of the Tokyo Metropolitan area, demonstrating how the slime mold forms strategic hub points that reflect real-world major population centers (i.e. Central Tokyo with the yellow nodes), akin to a real train system as shown in the adapted map of Tokyo railway^68^. **e**. A menu NetTracer3D offers to threshold the image by node degree, demonstrating one way network data can be joined to image analysis. The yellow highlighted nodes are being selected to stay in the image, due to having a degree of 4 or 5 in this case. **f.** A menu showing a variety of network analysis histograms NetTracer3D can generate using NetworkX algorithms. Here, we generate a histogram showing the distribution of shortest paths in the network, the number of edges each node must traverse to reach any other node^69^, a marker of network efficiency. **g**. A demonstration of NetTracer3D’s interactivity. Here, we select a node and can right click to reveal a menu with processing options. **h**. In g, we select ‘Show Neighboring Nodes and Edges’, resulting in the selected region to be expanded to encompass the aforementioned node’s neighbors. **i**. Demonstrating an additional neighbor search, now encompassing the neighbors of neighbors for our original node. Neighbor searches can be conducted from any number of nodes in this manner. **j**. Interacting with the network table. In this case, our selected subgraph from H is being displayed in the ‘Selection’ menu within the network table. We can save this selected region as a .CSV/.xlsx file to export it as a spreadsheet, or we have the option to save it in formats readable by other network graph analysis platforms, such as Gephi^70^. Source data provided.

### Communities and microenvironment detection

Beyond basic topological analysis, we include features where network nodes may be partitioned into communities using established community detection algorithms. There are three primary modalities offered.

1. The first utilizes **network community detection**, where densely connected subgraphs are partitioned due to having enhanced intra-group connectivity. Network community detection allows the separation of functional or spatially distinct clusters. The Louvain algorithm^19^ is implemented for modularity optimization, providing hierarchical community structure identification that has demonstrated efficacy in biological network analysis^20^. Additionally, label propagation algorithms were employed as an alternative approach, offering linear time complexity and parameter-free operation particularly suitable for large-scale network analysis^21^. Both approaches generate community-specific statistics including modularity scores—quantifying the strength of division into communities—and community-specific metrics that characterize the quality and biological relevance of detected partitions. Community partitioning can further be used to characterize the compositions of clusters of objects in the network, or just a way to simplify the number of objects into more macro-structures. For applications involving multiple node types, identified communities may be further aggregated into ‘supercommunities’ via K-means clustering^22,23^, joining functional network analysis with microenvironment characterization.
2. The second utilizes **neighborhood detection** via proximity networks. Proximity networks may be assembled by connecting to n-nearest neighbors, within a preset distance, or both. The immediate neighborhood for each node is quantified to assemble a set of vectors where each index describes the proportion of some node identity represented in that node’s neighborhood. Partitioning via K-means clustering is an established methodology to successfully detect neighborhoods in tissue^24^, but NetTracer3D allows a large amount of flexibility through the diverse parameters in establishing proximity networks.
3. The third offers **detection based on the channel intensity profile**, where the intensity of each node across the set of nodes is normalized, used to create a feature vector per node, which finally is clustered via K-means, a general pipeline that has been established for detecting significant, unique groups of cells^25^.

These approaches enable the identification of tissue microenvironments and cellular neighborhoods that recapitulate biologically meaningful functional organization patterns observed in complex tissues. The resulting neighborhood classification supports downstream analyses including component heatmaps and UMAP visualization, providing biologically pertinent descriptors of large-scale tissue domains and enabling quantitative characterization of cellular spatial relationships in microscopic images.

### Connectivity Networks — NetTracer3D Reveal 3D Neural Patterns in the FTUs of the Kidney and the Lung

Connectivity Networks establish inter-node connections via secondary objects identified within the edges channel, facilitating analysis of structural relationships between FTUs. This approach proves particularly valuable for evaluating connectivity patterns of FTUs, which are defined as the smallest tissue organization that performs a unique physiologic function and is replicated multiple times within an organ^4^. Such networks enable quantitative assessment of biological communication pathways, including neural and vascular connections that are critical for tissue function and disease progression^26,27^ where accurate mapping of these vascular-neural interactions has proven essential for clinical applications^28^. This functionality empowers NetTracer3D to evaluate complex relationships between separate FTUs, addressing a fundamental challenge in microscopy. This integrated approach is essential for advancing our understanding of renal neural networks, particularly given their involvement in sympathetic feedback cycles and heart disease^29^ while individual nephrons may function as integrated units with their own “central command” systems through juxtaglomerular and/or macula densa cells^30^.

We previously identified^31^ a unique 3D relationships between cortical kidney FTUs and nerves that explain organ-wide communications through central hub points distributed throughout the kidney. This extensive neural architecture provided the critical pathways necessary to transmit coordinated signals to all segments of a nephron and its neighbors, establishing a comprehensive communication network throughout the kidney cortex^32,33^.

Expanding on this initial qualitative application, we augmented NetTracer3D to deploy quantitative network analyses focused on neural connections between heterogenous structures, in this case being glomeruli and collecting ducts (Figure 2a). This capability addresses a critical need identified in recent three-dimensional kidney mapping studies, which have demonstrated the importance of comprehensive neurovascular connectivity analysis across the entire organ^34^. Our boundary-robust implementation of the Ripley’s Clustering Algorithm^13^ was then used to show both cell clustering behavior within glomeruli, then glomerular distribution in the image itself (Figure 2b), showing a relatively random distribution of cells within glomeruli, but a higher than random distribution of glomerular clustering within this area of the cortex.

**Figure 2.**
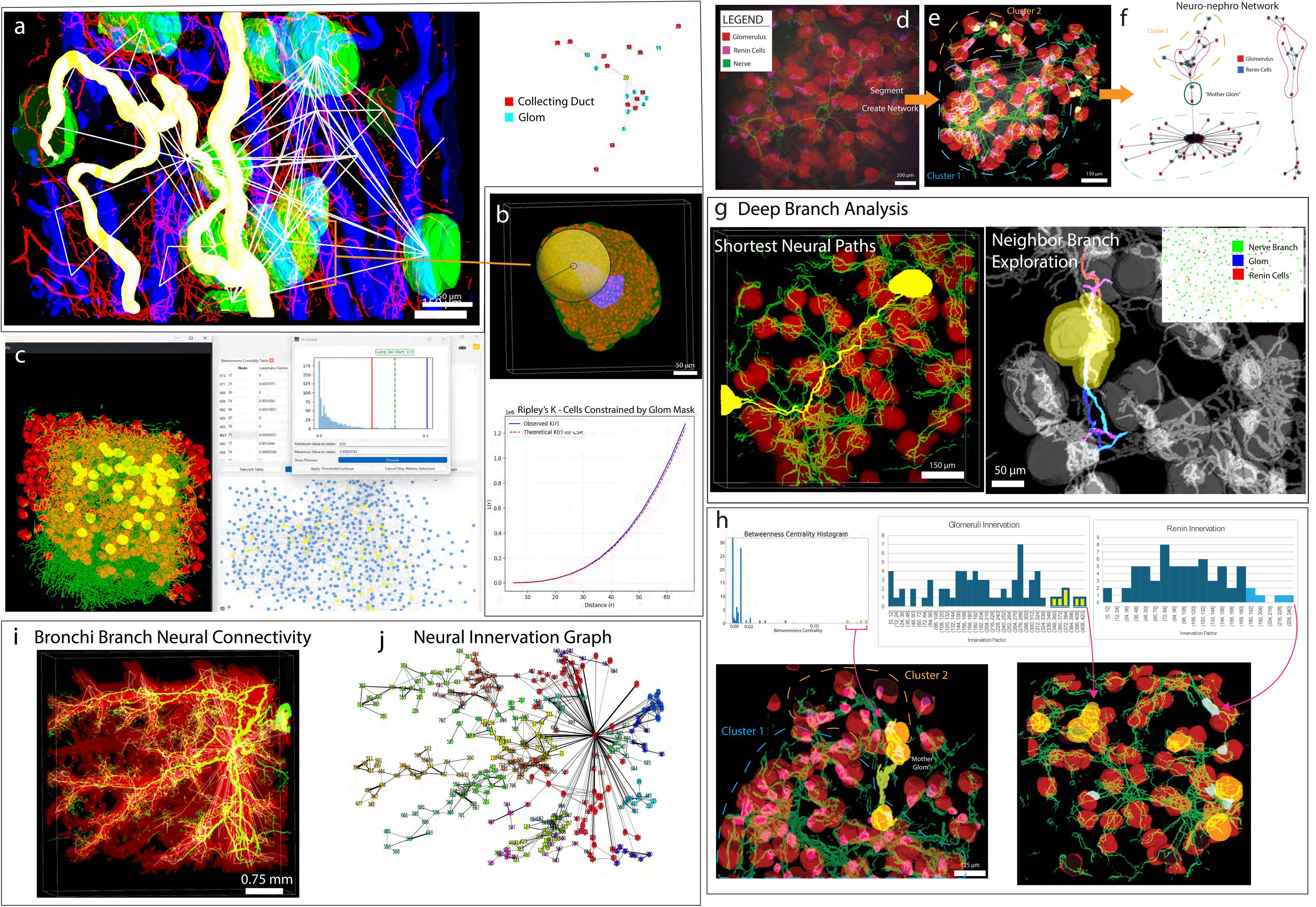
NetTracer3D Reveals Neural Connectivity Networks in Kidney and Bronchus of the Lung. **a**, (Left), Segmented collecting duct (blue), glomerulus (green), and nerve (red). The neural connections between collecting ducts and glomeruli are shown by the white lines. A larger plexus of merging collecting ducts (yellow) exists at the center of innervation, being downstream of multiple glomeruli-associated nerve projections into the medullary ray. (Right), The network graph of this structure, with collecting duct nodes colored red and glomerulus nodes colored cyan. The central duct is highlighted. **b**, One of the glomeruli, further used for a Ripley’s Clustering Analysis of its component cells. The glomerulus is shown in green, while its segmented DAPI-stained nuclei are red. To account for border artifacts, a safe search option enables search regions to only emanate from a constrained group of internal cells, shown in blue, while search regions themselves are kept entirely within the glomerulus mask, shown in yellow. The Ripleys’ K distribution is shown below. The red line represents a theoretical random arrangement of objects in the spatial boundaries, while the blue line represents the observed. In this instance it demonstrates a comparatively random layout. **c**, A view from NetTracer3D’s GUI. (Left), Segmented glomerulus (red) and nerves (green) from a 5x image of human kidney cortex. Yellow glomeruli have been selected from thresholding). (Top Right), Thresholding histogram to select gloms with high Betweenness Centrality. (Bottom Right), The integrated network graph also showing selected glomeruli (yellow). **d**, A 3D light sheet fluorescence microscopy image of mouse cortex with labeled glomeruli (red), renin cells cells (magenta), and nerve fibers (green). **e**, The same image, now segmented and transformed into a network. Two clusters of glomeruli in the network are circled in orange and blue, with the ‘mother glomeruli’ that joins them denoted in white. JGAs (denoted by the renin cells) participating in ‘JGA Neuro-Rosettes’ are highlighted yellow. **f**, The network graph with the aforementioned orange and blue clusters highlighted, with the mother glomeruli (and its JGA) circled in green. A third cluster that is not connected to the others within the field of view (FOV) was also observed. Red nodes are glomeruli, blue nodes are clusters of renin cells. The ‘JGA Neuro-Rosettes’ are circled in magenta. **g**, An example of exploring neural branches between nodes. (Left), The shortest nerve path between two distant glomeruli via nerve branches, all highlighted yellow. This function is not limited to branch analysis and can be used to quantify and visualize a variety of shortest path combinations. (Right), Labeled nerve branches were used to generate the network shown in the top right. In the 3D render, a selection of nerve branches off a glomerulus of interest have been isolated with branches denoted by the coloration schema. **h**, (Upper left), The betweenness centrality distribution of the networks’ nodes. (Upper Middle), The distribution of the magnitude of glomerular innervation. (Upper Right), The distribution of the magnitude of JGA innervation. (Lower Left), The nodes with the highest ‘betweenness centrality’ correspond to the mother glomeruli. They are highlighted in yellow alongside the nerves connecting the orange and blue clusters. (Lower Right), The glomeruli (highlighted yellow) and JG Cells (highlighted blue) that receive the highest magnitude of total innervation, within 10 μm. **i**, Image of a whole mount mouse lung with segmented bronchi and large vessel shown in red and sensory nerves in green. White lines represent neural connections between individual bronchial branch segments. Branches lacking innervation can be seen in the back. **j**, The network graph of the bronchial branch neural network. Neural connectivity can be observed to branch out from the central trunk of the bronchi, spreading along its branches. The branches participating in the neural network have been grouped into communities via Louvain clustering, denoted by node color. Source data provided.

A major discovery in our light sheet study was the existence of ‘Mother Glomeruli’, nephrons that enable inter-community communication between vascular trees of the cortex via the neural network. Employing NetTracer3D quantifications at scale, Figure 2c shows the use ‘Betweenness Centrality’^35^, one of the several network histograms that may be created, to algorithmically detect ‘Mother Glomeruli’ in a 3D volume containing 1000+ nephrons. This demonstrates how network results can be combined with more direct measures of spatial distribution within NetTracer3D, the only available software capable of describing these relationships.

As another demonstration of these multi-structure networks, we evaluated the relationship between the juxtaglomerular apparatus as a command center and corresponding glomeruli and their communities (Figures 2d**, 2e**). NetTracer3D was used to generate networks between glomeruli and nearby renin-expressing cells in the juxtaglomerular apparatus (JGA) of mouse cortex, a region that is crucial for blood flow to the glomerulus, blood pressure regulation and responding to changes in fluid status. This demonstrated the detection of mother glomeruli within mice (Figure 2f), demonstrating a conserved pattern of innervation between species.

NetTracer further allows analysis at high granularity that combines branch architecture with functional involvement of nodes within the network. For example, Figure 2g shows how branch labeling can be combined with connectivity networks to characterize connectivity through individual nerve branches and to detect shortest paths between structures in 3D space. Betweenness Centrality can be used to isolate these objects at high resolution (Figure 2h), along with other forms of analysis.

The connectivity networks offer the capability where virtually any structure that can be segmented into nodes (such as cells) can be connected via some additional path. To demonstrate these unique capabilities to other organ systems, we applied these ‘connectivity networks’ to investigate relationships between a primary branched structure (say vessels, or bronchi of lung) and a second connector (nerves, in this case). Through branch labeling, nodes may be yielded from any branched structure, whose connectivity via nerves are shown in Figs 2i, 2j. This analysis revealed that the nerves branched in a similar pattern as the bronchi themselves without evidence of notable downstream connectivity between bronchial branches. Interestingly, not all bronchi received innervation. Nerves initially tracked along all larger branches, however they were inconsistent about how far they would follow them. This was shown when many bronchioles received nerves from their parent bronchi, while those downstream of shorter nerves appeared to lack innervation. Differential innervation of bronchi may hold clinical implications as common medications like asthma nebulizers utilize similar mechanisms to sympathetic nervous signaling to function^36^. Additional examples of analysis on these datasets are provided in **ED** Figure 1. Overall, NetTracer3D enables a new class of quantitative 3D network analysis that was previously inaccessible with existing tools, with broad applicability demonstrated across tissues, from cellular aggregates such as the JG cells to large functional tissue units such as nephron segments or the lung bronchi.

### Branch Networks — Analysis of Cranial Vessels and Lymph Node

In support of diverse analytical applications, NetTracer3D incorporates sophisticated branch labeling algorithms designed to automatically decompose complex branched structures, such as neural networks and vascular trees, into their constituent branch segments^37,38^, building upon established methodologies for automated blood vessel segmentation and quantification in microscopic imaging. Each identified branch receives a unique label, enabling comprehensive structural analysis through multiple network paradigms. When integrated with Proximity Network algorithms, branch-to-branch networks can be constructed to analyze inter-branch relationships and connectivity patterns. Alternatively, Connectivity Network methodologies can be employed to investigate branch interactions with surrounding anatomical structures, providing insights into complex tissue architecture and functional relationships^39^.

We demonstrate NetTracer3D’s unsupervised approach to branch analysis (e.g., for neurovasculature), beginning with a magnetic resonance angiogram and proceeding through vessel segmentation to produce a spatially-aware network graph (Figure 3a). Unlike supervised methods that require CNN models to be trained on a training set^40^ NetTracer3D operates without training data dependencies, depending instead on a pre-obtained segmentation, addressing the domain adaptation problem where applying models across different datasets typically results in segmentation accuracy dropping “sharply”^41^.

**Figure 3.**
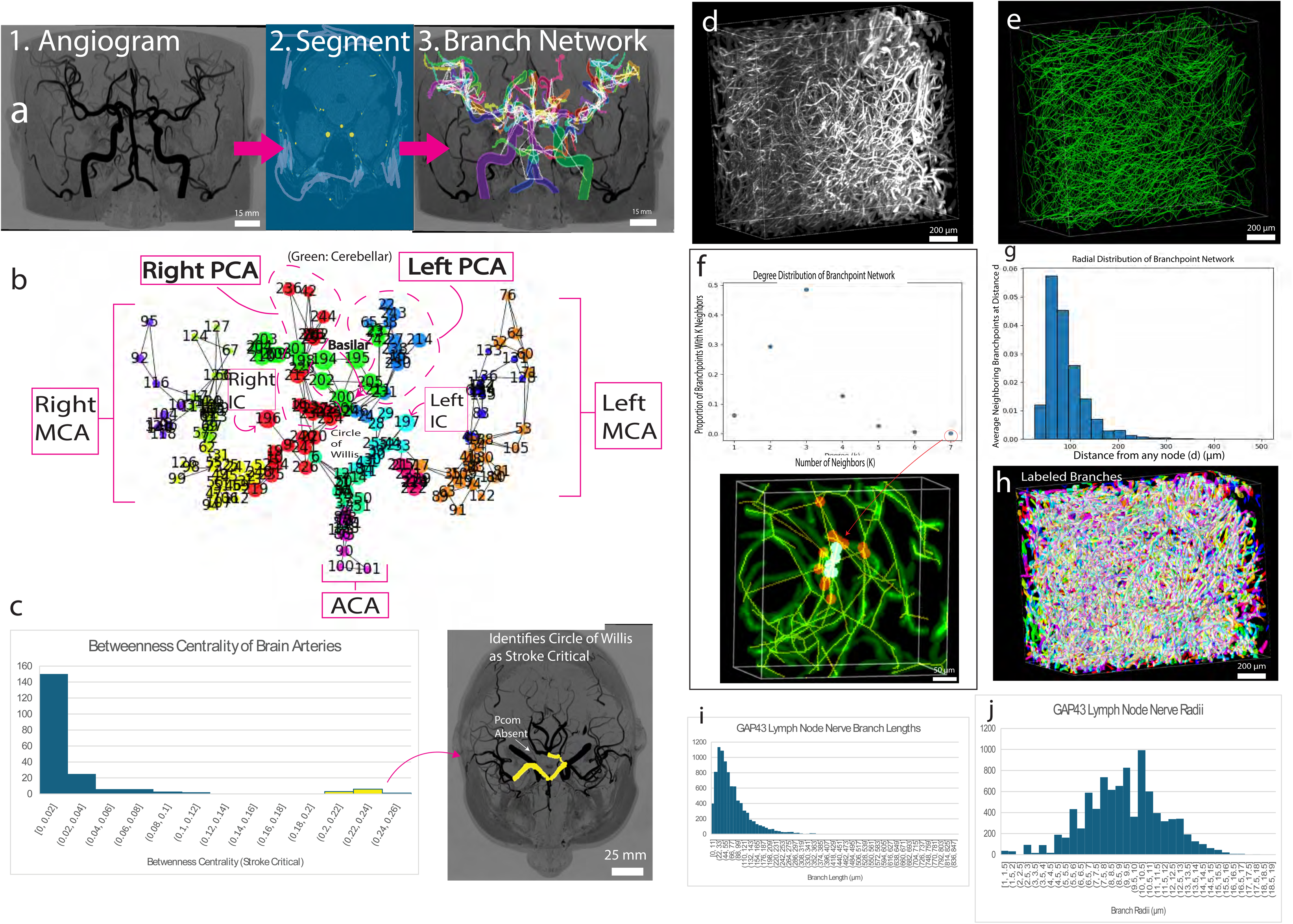
Branch networks of central arteries in the human brain and of neural patterns in a human lymph node. **a** (*Left)*, a Magnetic Resonance Angiogram showing blood vessels in the human brain. In the center, an example of vessel segmentation in NetTracer3D. *Right*, The segmented branches are labeled and connected in a network. **b**, The network graph of the above branch network. Vessels are placed in the graph reflecting their spatial location. Nodes are color-coded based on Areas of anatomical interest have been denoted. (MCA: Middle Cerebral Artery; PCA: Posterior Cerebral Artery; ACA: Anterior Cerebral Artery; IC: Internal Carotid). **c**, An example of how network analysis can be applied to spatial data. Here, we calculate the betweenness centrality^71^], showing nodes that represent crucial transit points in the network. A group of high-scoring objects matches almost exactly to the Circle of Willis, which in theory denotes these regions as highly disruptive stroke targets. **d**, A light sheet fluorescence microscopy image of Anti-GAP43 immunostained nerves from a 21-year-old female patient’s lymph node. **e**, The 3D visualization of extracted ‘branchpoint network’ of the lymph node, where adjacent branchpoints have been joined in a network graph. **f**, (Top), The degree distribution of the branchpoint network, demonstrating a mean degree of 2.8, a mode of 3, a maximum of 7. **g**, The radial distribution of the branchpoint network. (Bottom), One of the branchpoints with a degree of 7, highlighted in blue. Its neighboring branchpoints are highlighted in red, overlayed on a selection of the original lymphatic image (green). Connections between branchpoints are shown in yellow. **h**, The labeled branches derived from the segmented lymphatic vessels. **i**, The distribution of branch lengths of the aforementioned lymphatic nerves, as calculated by NetTracer3D. **j**, The distribution of radii of the lymphatic nerves. Source data provided.

The resulting network graph (Figure 3b) preserves spatial relationships while enabling network analysis. Vessels are positioned according to their anatomical locations, with nodes color-coded to reflect different regions of interest. This spatial embedding addresses limitations in existing approaches that struggle to maintain both topological connectivity and anatomical relevance.

We also showcase NetTracer3D’s community detection capabilities (Figure 3b). After graph creation, the Louvain community partition algorithm was able to identify anatomically meaningful vessel groupings corresponding to major cerebrovascular territories (MCA, PCA, ACA, IC) without supervision. This addresses a critical gap in automated vascular analysis, as traditional methods either require extensive manual annotation or fail to capture anatomical organization. The community detection successfully clusters vessels into regions that align with established neuroanatomical knowledge, demonstrating the method’s ability to discover clinically relevant structures without prior training. While such unsupervised clustering will not guarantee 1-to-1 correspondence with conventional anatomical grouping, they offer significant advantage in terms of speed, handling biological variants, and exploratory datasets, such as neovascularization that lacks formal grouping conventions.

As an example of downstream analysis, Figure 3c represents a novel application of using the betweenness centrality analysis of the network to identify critical vascular branches. Taking advantage of NetTracer3D’s interactivity, we identify high-scoring areas and utilize option to select groups of nodes, highlight them, and display the selection in 3D with Napari^17^. This analysis yielded the Circle of Willis region, aligning with clinical knowledge that “the arterial connections in the Circle of Willis are a central source of collateral blood flow and play an important role in pathologies such as stroke and mental illness”^42^. The Circle of Willis is an important area of anastomosis in the human brain, with abnormalities to this region associated with ischemia-risk^43^. For this particular image, one posterior communicating artery was absent from the initial image, a common biological variant^44^, and therefore is the only portion of the circle not appearing. The high centrality scores in this region nonetheless reflected its importance as a transit hub in the cerebrovascular network, providing automated identification of structures critical for stroke risk assessment.

NetTracer3D operates without GPU requirements, making advanced cerebrovascular network analysis accessible without advanced equipment. The integration with NetworkX provides access to established graph algorithms while maintaining the spatial context essential for meaningful anatomical interpretation.

To further enhance analysis of complex branched biological structures, a second branch-analysis implementation within NetTracer3D focused on creating “branchpoint networks”, a computational framework that transforms arborizing structures such as neural networks and vascular systems into quantifiable network representations of interconnected branchpoints. This analytical methodology enables precise quantification of branching, tortuous structures that have historically been challenging to analyze systematically^45,46^. This multi-scale network analysis capability supports investigation of both local and global organizational principles governing biological systems, from microscopic vascular architectures to tissue-level functional connectivity patterns^47^ enabling comprehensive characterization of complex biological networks across multiple spatial scales.

To demonstrate the clinical relevance and analytical power of this methodology, we present our network analysis derived from immunostaining for Anti-GAP43 (a critical axonal growth cone protein^48^) in a human lymph node obtained^49^ (Figure 3d). These lymphoid neural networks have been shown to locally modulate gene expression patterns in diverse cellular populations, including endothelial cells, stromal fibroblasts, and various leukocyte subsets, thereby highlighting their critical neuromodulatory potential in immune surveillance and response coordination^50^.

Previous investigations have documented that GAP43-positive nerve fibers within lymph nodes undergo dynamic sprouting and spatial redistribution into red pulp regions, anatomical compartments where immune cell populations predominantly reside, following immunization compared to control conditions^51^. This neuroplasticity response suggests that neural remodeling may play a crucial role in adaptive immune responses and lymph node functional reorganization. Figure 3d demonstrates the raw three-dimensional LSFM image capturing the intricate neural architecture within the lymphoid tissue. Figure 3e subsequently presents the internal network structure extracted through NetTracer3D processing, transforming the complex three-dimensional neural morphology into an easily quantifiable topological representation that preserves essential connectivity information while enabling systematic statistical analysis. The direct analytical outputs from this computational approach are illustrated in Figures 3f and **3g** which provide quantitative insights into neural network organization. Figure 3f displays the comprehensive degree distribution analysis, which quantifies the neighbor connectivity for each branchpoint within the network topology, a direct measure of local “branchiness” or structural complexity^52^. Our measurements revealed that the modal number of new branches emerging from any given branchpoint was 3, with a mean degree of 2.8 and a maximum branching degree of 7, indicating right-skewed heterogeneity in local neural complexity. This has direct application to a broad variety of biological analysis of innervated structures, quantifying the arborization of any neural structure, a characteristic that may be up-or-downregulated in various neural pathologies^53^. NetTracer3D enables detailed examination of specific structures of interest through targeted analysis capabilities. Following extraction of the complete degree distribution table, we searched for nodes exhibiting maximum connectivity (degree = 7), directed NetTracer3D to identify and highlight neighboring nodes, performed targeted cropping of the surrounding anatomical region, and visualized the results using the Napari^17^ software. This focused analysis revealed the presence of a complex neural plexus within the lymph node architecture, displayed in cyan coloration, with neighboring branchpoints highlighted in red (Figure 3f**, bottom)**. This analytical functionality enables rapid isolation and detailed examination of any high-throughput region of interest, facilitating both discovery-based research and hypothesis-driven investigation of specific anatomical features. Figure 3g presents the radial distribution analysis, which characterizes the spatial distances required to encounter subsequent branching events, providing insights into the neural network’s spatial organization and density patterns^54^.This metric offers valuable information about the efficiency of neural coverage within the lymphoid microenvironment. The labeled branches themselves (Figure 3h) can be used to obtain various direct morphological measurements, such as branch lengths (Figure 3i) and radii (Figure 3j). Additional examples of branched analysis, including reunification of large major branches beyond branchpoints and a demonstration of the filament tracer, are shown in **ED** Figure 2.

### Proximity Networks – Characterizing the 3D Tumor Microenvironment using CODEX and Analysis of 3D Stimulated Raman Spectroscopy Data

Proximity Networks represent an alternative network construction approach based on distance-based relationships, utilizing either Euclidean distances between node boundaries or centroid-to-centroid distances computed through efficient KD-tree clustering algorithms^55^ which provide scalable spatial partitioning for high-dimensional datasets. This methodology enables network generation from centroid coordinates alone, eliminating the requirement for complete image datasets. This capability is advantageous when processing extremely large node populations or integrating downstream analysis results from established pathology software platforms such as QuPath^56,57^ which provides comprehensive open-source bioimage analysis capabilities for whole slide image analysis. To facilitate seamless software-to-software integration workflows, NetTracer3D incorporates a secondary dedicated graphical user interface specifically designed for efficient data transfer from CSV spreadsheets to Network3D objects, enhancing interoperability within existing computational pathology ecosystems.

To demonstrate a proximity network application in the setting of a large number of channels and cells, we used a multiplexed dataset of a human tonsil, imaged using CODEX (Figure 4). Analysis begins with the segmentation of cells to create a mask where each bears a unique numerical label. A major challenge in multiplexed imaging is assigning cell phenotypes based on marker expression. Marker intensities can vary over several orders of magnitude, preventing universal cutoffs for marker positivity, while tissues often exhibit background staining from autofluorescence or non-specific binding. Additionally, unsupervised clustering followed by manual annotation requires subjective choices and can reduce labeling accuracy with increasing cell-type granularity^58^.

**Figure 4.**
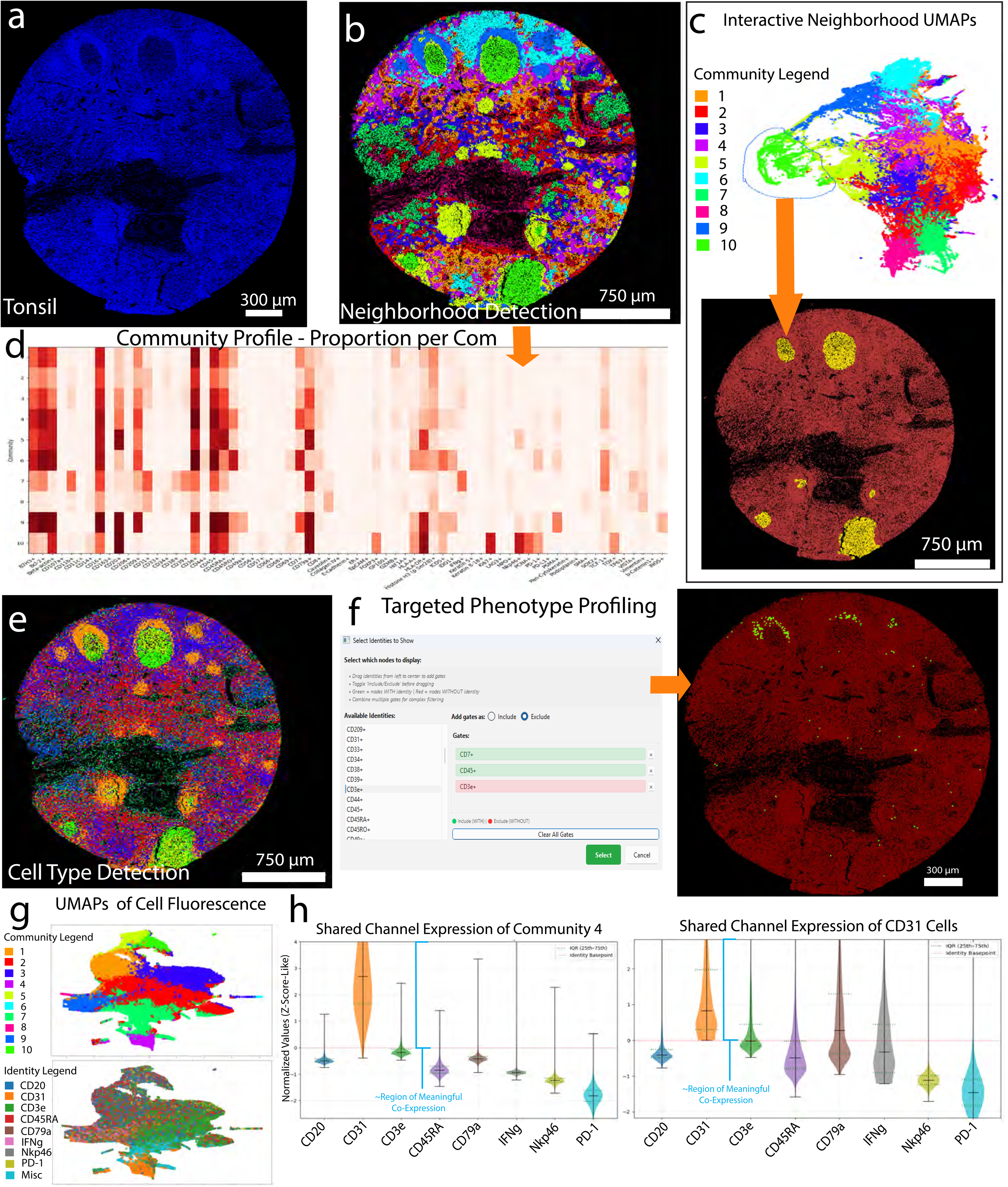
Proximity Network Analysis of a CODEX image. **a**, 2D CODEX Human Tonsil DAPI Channel. **b**, (Left), The result of partitioning the cells into 10 communities based on the different expression of markers within each neighborhood, with marker expression being defined by user threshold gating. Note this can be performed directly on the data without user gating, however we found this led to more loosely defined communities. (Right), Histologically significant regions such as germinal centers (neon green) can be extracted with this method of analysis. The mantle (yellow) is clearly visible as well. **c**, (Top), A UMAP showing the relative differences of our communities (colors and legend matching the previous figures), with further regions being appraised as ‘more different’. The UMAP is interactable, with the blue circle representing how a user would select a particular region. (Bottom), This selection is automatically mapped back to the segmented image in the image viewer window, as shown by the highlighted regions where the user in this case has selected the germinal center community on the UMAP. Selected objects may be manually assigned to their own communities, allowing for flexible UMAP partitioning as needed. **d**, Proportional breakdown of each expressed marker across 67 markers for our 10 communities. **e**, The result of partitioning the cells into 10 communities based on the normalized intensities across all channels for each node. This method does not detect smooth regions like **4b**, but rather, individual classes of cells, as shown by the increased partitioning of the germinal centers. **f**, Example detection of specific parent classes of cells from multiple identities. (Left), We select CD45+, CD7+, CD3e-cells as a representation of natural killer cells. (Right), The found cells (green) are shown to be in the paracortex immediately above the germinal centers. For more tailored cell discovery, combinations of the presence or absence of markers may be utilized to assign each cell a score, which through thresholding can be used to select groups of cells resembling a desired phenotype, without relying on absolute identity gating. **g**, (Top), A UMAP showing the relative differences of our communities (colors and legend matching **Figure 4e**), with further regions being appraised as ‘more different’. (Bottom), The same UMAP, but now labeled by a selection of node identities. Node identity colors are separate from previously shown community colors. **h**, (Left), Violin plots showing normalized expression of channel intensities for community 4 from **Figure 4e**). Areas above the red line represent ‘valid expression’ as defined by the lower bound of each marker by the user. In this case, community 4 largely represents endothelial cells, denoted by the strong expression of CD31 over other markers. In our upper UMAP in **Figure 4g** we can see community 4 (magenta) towards the bottom, while in **Figure 4e**, it may be observed lining blood vessel walls. In our lower UMAP, we may observe how community 4 largely overlaps with CD31-containing regions. (Right), Violin plots showing normalized expression of channel intensities for marker CD31. Here, we see a small amount of overlap with our CD31 cells and the upper quartiles of immune markers CD3e and CD79e. These likely represent the presence of these immune cells in immediate proximity of endothelial cells, ie in the process of egress, rather than true overlap.

To address these limitations, NetTracer3D implements an interactive thresholding approach that maintains direct cell ID integrity throughout the analysis pipeline. After initial Hoechst or DAPI-based segmentation (Figure 4a), NetTracer3D calculates mean marker expression for each cell (with optional cytoplasmic expansion) and presents real-time threshold histograms based on actual cell intensity distributions. Users can define both upper and lower expression bounds for each marker, enabling precise control over background noise and non-specific staining while preventing classification errors from signal overlap. This directly addresses the well-documented issue of spillover between neighboring cells that causes false positive co-expression of mutually exclusive markers^59^. By keeping users actively involved in the thresholding process, NetTracer3D provides a transparent alternative to black-box approaches that rely on unpredictable automated or pretrained machine-learning algorithms. This user-in-the-loop solution enables cellular phenotyping across numerous channels requiring only a single segmentation easily obtained from software such as Cellpose^60^. To replicate results between similar sets of tissues, the user may optionally train a neural network to replicate their minimum and maximum threshold preferences, allowing automation of the thresholding process based on their unique datasets. If quick analyses is desired, cell identity assignment may be skipped to solely cluster based on neighborhood Z-score intensity profiles, although this loses access to some downstream features that make use of cells having a binary identity state. Finally, artificial hexagonal or rhombic dodecahedrons may be generated to serve as nodes if a cell segmentation is not available, providing neighborhood analysis within any spatial domain.

Figure 4b shows the results of this neighborhood clustering pipeline, yielding clearly identifiable regions consistent with lymphoid follicles in the tonsil. This includes the germinal center (neon green), mantle (yellow rim around germinal center), paracortical areas (sky blue, light blue). Blood vessel associated regions are detected by the pink and red communities, along with other communities. The results of the K-Means clustering was cross-verified by UMAP visualization (Figure 4c). These UMAPs generated by NetTracer3D are fully interactable, allowing the user to freely circle, select, and manually partition regions of the map. Finally, the compositional breakdown of each community is represented by the heatmap in Figure 4d.

We also demonstrate the results of community clustering based on the Z-score fluorescent intensity profile to detect individual, unique classes of cells rather than communities (Figure 4e). The results are less ‘smoothed-over’ than Figure 4b, more prone to selecting unique cells despite their neighborhood composition, as shown by splitting of the germinal centers into two cell classes with a few others scattered within, consistent with established divisions between light and dark zones within the germinal centers. This strategy can be used to discover distinct cellular identities and phenotype combinations, which themselves can be combined to pick up unique cell types. For example, Figure 4f shows the combination of CD45+, CD7+, and CD3e-as a way to identify natural killer cells^61^, found here to be abundant in the paracortex above each germinal center.

In addition to verifying K-Mean’s clustering results, UMAPs can be used to depict cell identities. In this case, 8 of our 67 identities are plotted for ease of visualization, with cells lacking those markers assigned to the ‘misc’ category (Figure 4g). The relationships between these markers are expanded on in NetTracer3D’s violin plots (Figure 4h). If the user had previously assigned cell identities via thresholding, these violin plots are normalized so that 0 represents the lowest example of what they consider a ‘true’ instance of that identity. In short, any violin plot region above 0 equates to true co-expression of that channel, as defined by the user.

These violin plots may be alternatively used to display the amount of ‘valid’ expression of each channel for any community/neighborhood. In this instance, we explore the expression profile of community 4, as well as the CD31 endothelial marker. We observe that community 4 appears to select endothelial cells on Figure 4e; correspondingly, the leftmost violin plots demonstrate strong CD31 expression in community 4 with little expression of the other sample markers. When we evaluate CD31’s shared expression with other markers throughout all nodes in the rightmost violin plots, we now see some level of intensity overlap with certain immune markers such as CD3e; this may represent CD31 expression which has been previously noted in certain thymocytes.

This pipeline can be seamlessly applied to 3D tissue slices, allowing characterization of multichannel objects and neighborhoods throughout a 2.5D (i.e., serially sectioned) or a true volumetric 3D image. In Figure 5, we characterize a 3D Stimulated-Raman Spectroscopy (SRS) slice of human kidney (Figure 5a**, b)**, a form of imaging that uses label free infrared spectra to characterize tissue analytes such as protein, lipid saturation and unsaturation, and metabolites^62^. To standardize intensity-based appraisal of nodes through the Z-stack (where light may become dimmer in the deeper tissue), NetTracer3D offers a normalization algorithm that selectively brightens deeper tissue regions based on a globally derived change in intensity vector field, maintaining local differences in brightness, applied in Figure 5b. We then generate hexagonal prisms with a height and side-length of 8 microns, each extending through two slices of the Z stack. Since we lack a cell channel, NetTracer3D offers a simple workaround suitable for 3D data via the generation of artificial nodes (Figure 5c**, d)** from either hexagonal prisms or rhombic dodecahedrons to achieve ideal tessellation^63^. Similar to the tonsil example, these nodes acquire a metabolite ‘identity’ (Figure 5e) via manual thresholding. Like before we can use nearest-neighbor based K-means clustering, but this time, nearest neighbors are detected in Z as well as the X/Y plane, resulting in three-dimensional neighborhoods (Figure 5f). The nodes themselves may also be clustered based on their metabolite Z-scores (Figure 5g). In Figure 4d, we demonstrated how these neighborhoods may be described by their proportions. Figure 5h now shows how they may be alternatively described by ‘enrichment’, that is, the ratio of cells in a neighborhood versus outside of it. The former is ideal for characterizing the tissue type by its major components, while the latter allows detection of where expression levels are abnormally high and where rare markers tend to appear.

**Figure 5.**
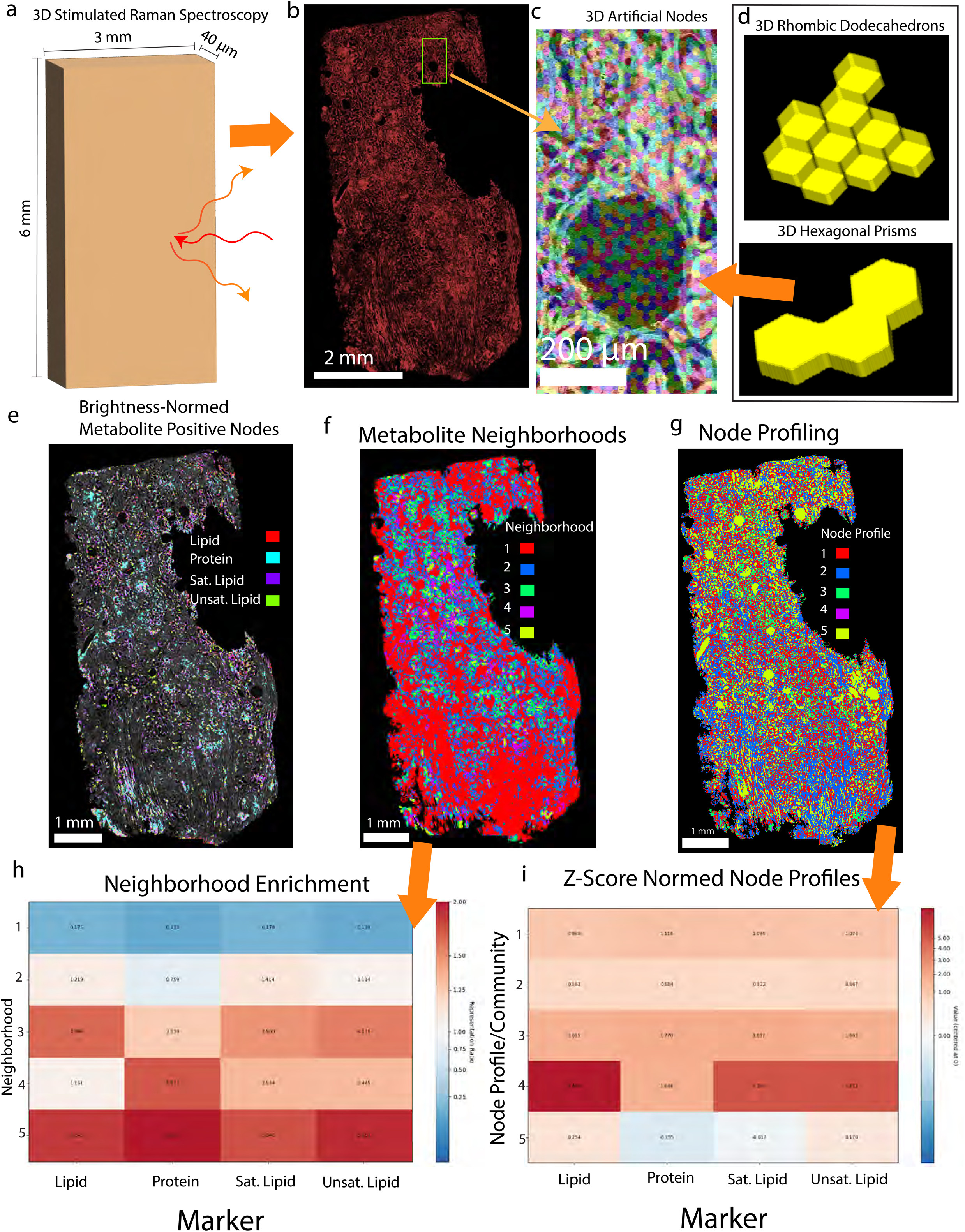
Analysis of 3D Stimulated-Raman-Spectroscopy Slice with Proximity Network. **a**, A 3D graphic of our tissue slice showing its dimensions. The slice is rather tall and wide, while being 40 microns thick, enough to pick up several layers of cells. **b**, A picture of the protein channel from this image. This tissue is a thin 3D-slice containing primarily human kidney cortex and some medulla towards the bottom. **c**, A closeup of the protein channel (gray) overlayed with the 3D hexagonal prisms (multicolored) that serve as artificial nodes. **d**, The actual 3D appearance of the (Top) rhombic dodecahedrons and (Bottom) hexagonal prisms that NetTracer3D generates to serve as artificial 3D nodes. For 2D data, hexagons will be used by default. **e.** A slice from the tissue showing which nodes have now picked up specific metabolite identities (as denoted by the legend). **f**, The same slice after neighborhood clustering is applied to create 5 metabolite-based neighborhoods. **g**, Once again, this slice, now after channel-intensity based clustering is applied on the nodes themselves to group similar nodes into 5 communities. **h**, The enrichment profile describing **Figure 5f**. Values greater than 1 represent relative enrichment while values less than 1 represent relative deficiency of a marker. Neighborhood 1 can be observed to be a relatively low-expression neighborhood. **i**, The Z-score profile of individual nodes describing **Figure 5g**. In this instance, nodes of community 5 were also relatively under-expressive, mostly picking up nodes in acellular areas such as the center of tubules or spaces where glomeruli had physically fallen out of their Bowman’s capsules (leaving a cavity).

As 3D imaging capabilities have advanced through breakthroughs in imaging and clearing technologies^64^, NetTracer3D fills a crucial gap in accessible analysis of 3D cellular data. We lastly demonstrate NetTracer3D’s suitability for large-3D object analysis with a tumor-stroma spheroid multiculture from Diosdi et al^65^ (Figure 6a**, b**). After generating a proximity network, we opt to detect spatial clusters via network community detection algorithms like Louvain clustering^19^ (Figure 6c). These communities may be further clustered into supercommunities, or have their compositions described as shown in Figure 6d. Here, we detect three clusters that contain overlap between Fibroblast, Endothelial, and/or Tumor markers. The fluorescent profile of two of these clusters are shown in Figure 6e, where violin plots were generated showing differing valid intensity expression levels for each channel within any community. Additional 3D spatial analysis without network requirements may be performed, such as generating spatial heatmaps based on clustering behavior with flexible parameters (Figure 6f), where clustering color is based on theoretical distribution throughout the 3D convex hull of the binary dataset. In this case, we generate a nearest neighbor heatmap of the tumor cells relative to distance to fibroblasts, confirming the presence of increased proximity of those cell types in the center of the spheroid, as verified by the composition of the central community 3. Finally, we demonstrate how nearest neighbor distances may be calculated and visualized within a graphical matrix for all identity combinations (Figure 6g).

**Figure 6.**
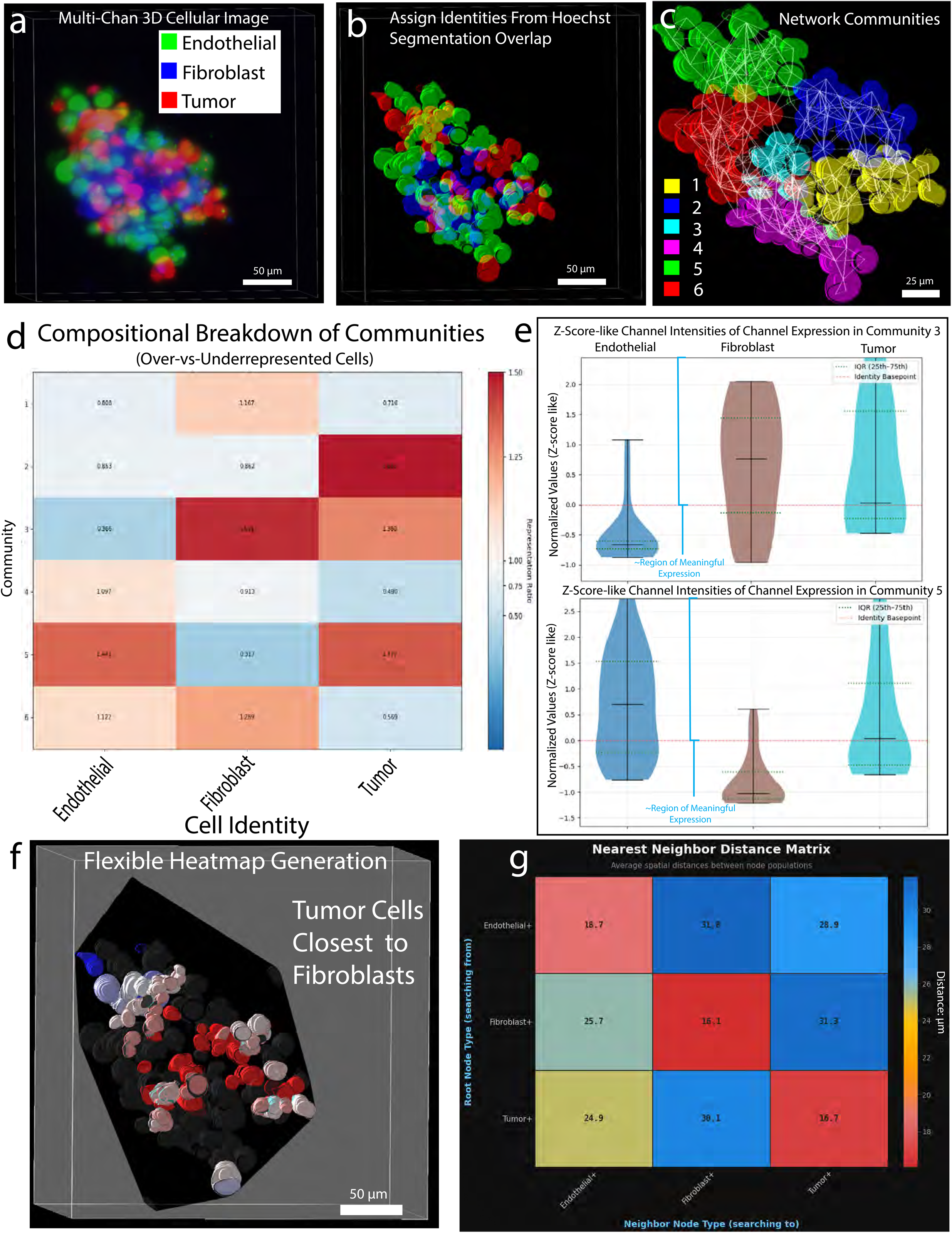
Characterization of 3D Tumor Microenvironment with Proximity Network. **a**, Analysis of tumor-stroma spheroid multiculture from study by Diosdi et al^65^. Multiculture 3D tumor spheroids mimic solid tumors in many aspects, such as the heterogeneous architecture, growth kinetics, physical interactions or complex communication. Channel stain schematic: (Nuclei – ch001, EA.hy926 – ch002 [Green – Endothelial Cell Line], tumor – ch003 [Red], and MRC-5 – ch004 [Blue – Fibroblast Cell Line]). **b**, Solid ellipsoids represent individual cells after segmentation of the Hoechst nuclear channel. Colors indicate identity assignment based on mean intensity overlap in each channel combined with human-in-the-loop thresholding. **c**, Example proximity network. Cells connections were formed within 10 µm with search from node borders rather than centroids. In this case, cells were not permitted to form connections over other neighbors’ search regions, which served to aid in logical spatial partitioning. Cell coloring denotes which network community each belongs to. **d**, Heatmap graph showing the over-vs-underrepresentation of cell types within each community relative to the abundance of those cells in the entire image. > 1 (red) means overrepresented; < 1 (blue) means underrepresented. This was derived from heatmaps pertaining to community composition, which may also be generated. **e**, Violin plots charting the distribution of fluorescent marker expressions for cells in communities number 3 (top; cyan in **Figure 4c**), a centrally located community expressing fibroblasts and tumor markers, and number 5 (bottom; green in **Figure 4c**), a peripherally located community expressing fibroblast and endothelial markers. Regions above 0 represent ‘true expression’ of that marker within the cells of the community, based on the lower threshold bound assigned by the user during cellular identity assignment (shown by the red dotted line at 0). Expression values above and below 0 are transposed, then normalized based on their Z-Scores. It can be observed how these values based on ground truth fluorescence mirror the community representations in **Figure 4d**. **f**, Example of spatial heatmap generation. Heatmap colors are assigned based on comparison with theoretical distribution where cells are evenly spaced, with red representing closer neighbors than expected, and blue representing more distant. This theoretical distribution may be confined to the convex hull of the 3D objects (shown in black), allowing theoretical comparison to occur without interference by the image background. In this instance, we opt heatmap generation to color tumor-expressing cells, with distances relative to the nearest 5 fibroblast cells, confirming the presence of increased proximity towards the center where we discovered community 3. **g**, NetTracer3D allows batch computation of nearest-neighbor relationships between all unique identities. It will return the chart shown, color coded with red representing the nearest neighbors and blue representing the most distant. This allows for rapid visual appraisal of the spatial nearness of any number of identities of interest. For more analysis more tailored to characterizing cell interaction, this method can also be employed to generate a similar chart that quantifies presence of network neighbors between cell cohorts, allowing bordering cells to be quantified based on a proximity network within a desired distance.

#### Morphological Analysis and Visualization

In addition to its network generation functionalities, NetTracer3D provides options for morphological analysis of structures, such as calculating and thresholding by radii, volumes of objects, non-network derived clustering analytics, and more.

#### Discussion

We developed NetTracer3D to provide an accessible user interface that streamlines methodologically complex pipelines and analyses of both 2D and 3D microscopic and medical images. The software delivers end-to-end pipelines encompassing image segmentation and processing, transformation to network data, and comprehensive analytical options for examining spatial, phenotypic, and morphological arrangements of objects. Our analysis identified three broadly applicable modalities for using NetTracer3D across diverse scientific imaging applications. First, the capabilities to discover **Connectivity networks** and enable analysis of functional tissue units (FTUs), cells, or other biologically relevant nodes connected via secondary structures such as nerves or vasculature. We demonstrated how this approach elucidated 3D neural connectivity among FTUs in human kidney cortex and revealed neural connectivity patterns among bronchial branches in mouse lung. Second, features to detect **Branch Adjacency and Branchpoint networks** facilitate the conversion of any branched structure into analyzable network representations. Given the prevalence of branched architectures in biological systems—including nervous, vascular, and lymphatic networks—this analytical framework has wide-ranging applicability from anatomical to microscopic scales including medical imaging datasets as demonstrated with analysis of human MRA cranial vessel datasets and nerve architecture within 3D LSFM images of lymph nodes. Third, **Proximity network** features provide a versatile approach for grouping structures based on spatial nearness. This modality proves particularly valuable for analyzing cellular organization; combined with community partitioning algorithms, we validate its utility by identifying biological meaningful clusters of distinct cell types within our CODEX lymph node, before providing more direct 3D analysis examples. With recent advances in multiplexed 3D imaging^9^, NetTracer3D addresses a critical need for accessible, user-friendly analysis tools in this rapidly evolving field. With the need for tools to characterize the tumor microenvironment from 3D images, such as those generated by the Human Tumor Atlas Network^66^, NetTracer3D’s suite of accessible, functional out-of-the-box analytical tools provides researchers an important resource.

Beyond network analysis, NetTracer3D provides an extensive toolkit for image segmentation, processing, and basic morphological characterization. Users can isolate and highlight specific objects of interest for targeted investigation. Through integration with napari software and the generation of customizable informative overlays, NetTracer3D facilitates robust 3D visualizations that are essential for in-depth qualitative analysis and the creation of publication-ready figures, including every figure in this paper. Collectively, these capabilities position NetTracer3D as a versatile platform that bridges the gap between image acquisition and biological insight across a wide spectrum of biomedical applications.

We acknowledge several limitations of NetTracer3D that present opportunities for future development. First, the tool’s reliance on segmented input data necessitates that users generate sufficiently high-quality segmentations, a requirement that may prove challenging for images with poor signal-to-noise ratios. Because NetTracer3D’s performance is sensitive to segmentation quality, we took deliberate care to include a variety of options to improve segmentations and screen out noise, such as our filament tracer or various forms of thresholding to remove noise artifacts. Second, many of NetTracer3D’s network generation algorithms include optional parameters that may require optimization to achieve ideal results for specific datasets. While default parameters are designed for broad applicability, we recommend independent verification and iterative adjustment until results meet the user’s quality standards. This consideration is particularly important for the branch labeling algorithm, especially regarding spine removal: each spine induces a branch split in the network representation. Finally, although NetTracer3D implements highly optimized Python algorithms leveraging packages such as NumPy and SciPy, the scale of certain 3D datasets, particularly LSFM images that can reach terabyte sizes, presents ongoing computational challenges that may necessitate downsampling to enable practical analysis.

NetTracer3D represents a significant advance in accessible quantitative analysis of 2D and 3D biological imaging data. As demonstrated throughout this work, these capabilities enable discovery of previously unappreciated biological phenomena with direct physiological implications: from the identification of ‘mother glomeruli’ that serve as critical neural hubs coordinating kidney homeostasis, to the characterization of anastomotic neural connections in lung bronchial networks, to automated detection of the Circle of Willis as a high-centrality vascular node relevant to stroke risk assessment, to spatial and phenotypic cellular characterization of CODEX images. Unlike existing commercial platforms such as Imaris and Amira lack comparable end-to-end network analysis pipelines, or open-source tools that require extensive coding knowledge, NetTracer3D and its GUI provide an accessible solution that allows general users to extract quantitative insights from complex spatial data across diverse tissue types and imaging modalities while remaining computationally lightweight on the CPU. NetTracer3D is freely available as an open-source Python package [https://pypi.org/project/nettracer3d/], and integrates seamlessly with established tools including napari^17^ for visualization and networkX for advanced graph analysis, positioning it as an immediately deployable resource for the broader biomedical imaging community. Collectively, NetTracer3D empowers users to generate and test hypotheses for use in research or medical applications/diagnostics as shown for loss of neuro-nephron connectivity in diabetic kidney disease^31^ or vascular abnormalities in radiological images.

## Methods

### Sample Collection

#### Mouse lung LSFM

##### Animals

Mice were euthanized in accordance with Institutional Animal Care and Use Committee (IACUC) guidelines and University of California, San Diego (UCSD) institutional policies. *Vglut2-IRES-Cre;Rosa26-tdTomato* mice were generated as previously reported^72^ to generate offspring expressing tdTomato in glutamatergic sensory neurons.

##### Tissue clearing and Immunostaining

Lungs from were *Vglut2-IRES-Cre;Rosa26-tdTomato* mice perfused with 10 ml cold PBS/heparin (100 µg/ml), placed into 15ml conical tubes with 4% PFA (PBS), fixed for 2Lhrs at 4°C, washed in PBS for 30Lminx2, and incubated overnight in PBS. Samples were stored at 4°C in PBS with 0.01% sodium azide until ready to process, up to 1 week. Tissue clearing was performed using iDISCO(ace) methods, as previously described^73^. Immunostaining was performed following initial sample dehydration, decolorization, and rehydration, with all steps carried out at room temperature (RT). Samples were transferred directly into blocking solution (0.2% PBST, 10% DMSO, 5% normal goat serum) and incubated overnight. Samples were then incubated for 2 days with primary antibodies diluted in PBS containing 0.1% Tween-20, 5% DMSO, and 5% normal goat serum: mouse anti-αSMA–FITC (1:600; Sigma, F3777) and rabbit anti-RFP (1:300; Rockland, RL600-401-379). Following primary incubation, samples were washed in PBS for 1 hr x 3. Next, samples were incubated overnight with secondary antibody diluted in PBS containing 0.1% Tween-20, 5% DMSO, and 5% normal goat serum: goat anti-rabbit Alexa Fluor 594 (1:600; Invitrogen, A32740). Lungs were washed in PBS for 2 hrs x 4, extended overnight, and then processed through the remaining clearing steps. For final clearing, samples were placed in ethyl cinnamate (ECi; Sigma, W2430000) and stored at RT until imaging.

##### Light sheet fluorescence microscopy

Lungs were imaged using a Zeiss Z.1 light-sheet microscope at the UCSD Microscopy Core facility. Agarose-embedded cleared lungs were suspended by puncturing agarose with a staple and suspending on a magnet. Samples were imaged in a large imaging chamber filled with ECi and light-sheet imaged with an EC Plan-Neofluar 5x (.16NA) Air objective. Lungs were imaged for sensory nerves (tdTomato, Channel 560/594) and pulmonary bronchi and vasculature (αSMA, Channel 488/FITC).

#### Mouse kidney LSFM

##### Animals

*Ren1c^+/−^*; *Ren1c-Cre*; *R26R^mTmG^*mouse (C57BL/6) was generated as previously reported^74^. In this mouse line, cells from the renin lineage become GFP positive upon Cre-mediated recombination while non-recombined cells express RFP^75^. It has been confirmed that *Ren1^c+/−^*mice are indistinguishable from wild-type mice in terms of blood pressure, urine volume, osmolality, kidney *renin* mRNA levels, the number of kidney cells producing renin, and plasma renin concentration^76^.

All animals were housed in a temperature– and humidity-controlled room under a 12-hour light/12-hour dark cycle. All procedures were performed in accordance with the National Institutes of Health guidelines for the care and use of laboratory animals and were approved by The Animal Care and Use Committee of the University of Virginia.

##### Tissue-clearing and whole-mount immunostaining

For the tissue-clearing and whole-mount immunostaining of mouse kidneys, we used the updated clear, unobstructed brain/body imaging cocktails and computational analysis (CUBIC) protocol^77^. Tissue-clearing and staining performed as previously described^78,79^. The mice were anesthetized with tribromoethanol (300 mg/kg) and perfused with 20 ml of phosphate-buffered saline (PBS) and 30 ml of 4% paraformaldehyde (PFA) via the left ventricle of the heart. Kidneys were removed, divided into two sections each, and fixed overnight in 4% PFA. Following all steps were performed with gentle shaking. The samples were washed with PBS for 6 hours. Then, the samples were immersed in CUBIC-L [mixture of 10 wt% N-butyldiethanolamine (MilliporeSigma, Burlington, MA) and 10 wt% Triton X-100 (MilliporeSigma, Burlington, MA)] at 37°C for 5 days. After the samples were washed with PBS for 6 hours, they were placed in blocking buffer (mixture of PBS, 1.0% bovine serum albumin, and 0.01% sodium azide) for overnight at room temperature (RT). Then the samples were immersed in immunostaining buffer (mixture of PBS, 0.5% Triton X-100, 0.25% bovine serum albumin, and 0.01% sodium azide) containing 1: 200 diluted Alexa647-conjugated anti-tubulin beta 3 (TUBB3) antibody (657406, BioLegend, San Diego, CA) for 16 days at RT. After washing again with PBS for 6 hours, the samples underwent post-fixation in 1% PFA for 3 hours, then they were immersed in 1:1 diluted CUBIC-R+ [T3741 (mixture of 45 wt% 2,3-dimethyl-1-phenyl-5-pryrazolone, 30 wt% nicotinamide and 5 wt% N-butyldiethanolamine), Tokyo Chemical Industry, Tokyo, Japan] at RT for overnight. The samples were then immersed in CUBIC-R+ at RT for 2 days.

##### Light-sheet fluorescence microscopy

Macroscopic whole-mount images were acquired with a light-sheet fluorescence microscope (Lightsheet7, Zeiss, Oberkochen, Germany). The samples imaged in mounting solution (RI:1.520) (M3294, Tokyo Chemical Industry) with Clr Plan-Neofluar 20×/1.0 Corr nd=1.53 detection optics. The voxel resolution was as follows: x = 0.472 µm, y = 0.472 µm, z = 1.00 µm (zoom × 0.5). GFP-expression was measured by excitation at 488 nm. TdTomato-expression was measured by excitation at 561 nm. The expression of TUBB3 was measured by excitation at 638 nm.

### Stimulated-Raman-Spectroscopy Kidney Slice

Fixed (4% PFA) kidney issue blocks were first embedded in 4% agarose solution before sectioning with a Comprestome (VF-310-0Z, Precisionary) to create 120μm-thick sections. These tissue sections were transferred to a glass slide with a 120μm thick spacer (Secure-Seal, 20mm diameter, Invitrogen), mounted with 1x PBS, and sealed with a number 1.5 thickness cover glass (Erie Scientific). 3D SRS methodology and acquisition parameters follow previously published protocols^80^ (https://dx.doi.org/10.17504/protocols.io.14egn9x2yl5d/v1), here applied for a whole-slide image at a depth of 40μm and for a 2mmx2mm ROI at a depth of 80μm. The acquired 3D image stacks were then processed utilizing a version of the VISTAmap algorithm (MATLAB) to remove the “vignette” effect found in stitched microscopy images^81^.

### Human Tonsil CODEX Image

Institutional Review Board approval (STUDY00012749) was obtained for the study of deidentified human tissue. A five micron histological section of formalin-fixed paraffin-embedded human tonsil tissue (2 millimeter circular punch) was examined by multiplex histology using a Phenocycler Fusion 2.0 (Quanterix) according to the manufacturer’s instructions. In brief, the sections were deparaffinized and rehydrated, followed by antigen retrieval with EDTA and citrate buffer. Autofluorescence removal was performed in a light bath (4.5% hydrogen peroxide, 27 mM sodium hydroxide in PBS) for 45 min. The slide was stained with the PhenoCode™ Discovery IO60 panel (Quanterix) of sixty DNA-barcoded antibodies, excluding antibodies against Keratin 14 and CD21. In addition, DNA-barcoded antibodies anti-HIF1alpha (Cat. 4550069, Quanterix) and anti-NKp46 (clone 29A1.4, gift of Sean Tracey and Michael Farrar, University of Minnesota) were included alongside the following antibodies which had been individually barcoded using an antibody conjugation kit (Cat. 7000009, Quanterix): Anti-CD45RA (Cat. No. 304102, Biolegend), anti-CD16 (Cat. No. 72204SF, Cell Signaling Technology), anti-CD7 (Cat. No. ab230834, Abcam), anti-CD49a (Cat. No. 328302, Biolegend), anti-CD69 (Cat. No. ab234512, Abcam), anti-Na+/K+ ATPase (Cat. No. ab167390, Abcam), anti-CD33 (Cat. No. MAB11371, R&D systems), anti-B7H3 (Cat. No. 58798SF, Cell Signaling Technologies) and anti-PSMA (Cat. No. 92728SF, Cell Signaling Technologies). Imaging was performed at 20x magnification on a PhenoImager (Quanterix). Complementary barcodes conjugated to Atto550, Alexa Fluor647 and Alexa Fluor 750 were applied alongside a DAPI stain in cyclical hybridization, buffer exchange, imaging and stripping cycles on the PhenoCycler (Quanterix). Image stitching, deconvolution, alignment based on DAPI, background removal and cycle concatenation were performed in Quanterix software to produce a 70-channel image.

## NetTracer3D Operations

### Network3D Object Properties

Important aspects of the dataset were stored as the Network3D class, containing the following properties:

**Nodes:** 3D labeled image array containing objects to be connected in network analysis, with each unique label representing a distinct node.

**Edges:** 3D labeled image array containing structures used to establish connections between nodes, such as filamentous networks or contact regions.

**Overlay 1 & 2:** Additional image channels used for visualization, temporary data storage, or intermediate processing steps.

**Network:** Graph structure storing node connectivity relationships, organized as node pairs with associated edge identifiers when available.

**Node Centroids:** Three-dimensional coordinates (Z,Y,X) representing the center of mass for each labeled node object.

**Edge Centroids:** Three-dimensional coordinates (Z,Y,X) representing the center of mass for each labeled edge object.

**Node Communities:** Community assignments for nodes derived from network clustering algorithms, grouping nodes with similar connectivity patterns.

**Node Identities:** Categorical labels assigned to nodes based on morphological, functional, or user-defined classification schemes.

**xy_scale:** Real-world dimension per pixel in the 2D xy-plane (e.g., microns per pixel), with equal scaling assumed for both x and y dimensions.

**z_scale:** Real-world dimension per voxel in the z-dimension, allowing for anisotropic scaling relative to the xy-plane.

### Main Window

The GUI was created using the Python PyQT6 module to serve as the graphics library. The main window employed three widgets. At the center, the ‘Image Viewer Window’ or ‘Canvas’ utilized matplotlib to generate 2D composites of all four channels, plus the highlight overlay. Cropping and image-pyramid style downsampling was used when possible to increase rendering time. The upper-right-widget (the tabulated data widget) was a specialized pandas dataframe that stored multiple tables. The lower-right widget (the network table) was another pandas dataframe that stored a table to show the network, and a similar table to show sub-selections within the network.

### Connectivity Network Calculation

Connectivity networks were calculated using a distance transform-based approach. Nodes were expanded based on a user-defined search distance using a distance transform that assigned outer shell regions labels corresponding to the nearest internal labeled node. The search regions were used to discretize edges into “inner edges” (within search regions) and “outer edges” (outside search regions), with each edge piece acquiring unique labels. For outer edges, single-voxel dilation was applied to ensure overlap with search regions. Inner edge connectivity was determined by extracting search region borders using the skimage find_boundaries method and identifying inner edge pieces within these borders through dilation-based contact detection. Network connections were established between nodes that interacted with the same labeled edge piece. The resulting node-edge relationships were converted to a NetworkX graph object for network analysis.

### Proximity Network Calculation

Proximity networks were calculated by connecting nodes within a user-defined distance threshold using two available algorithms. For centroid-based search, node centroids were normalized according to xy/z scaling parameters and connections were identified using the scipy.spatial KDTree data structure for efficient distance calculations between points. For morphological shape-based search, bounding boxes around labeled objects were identified using scipy.ndimage.find_objects(), and subarrays containing each node plus its search region were extracted. Node dilation was performed using scipy.ndimage.distance_transform_edt(), and the resulting dilated masks were applied to isolate neighboring nodes within the search distance. Node-neighbor relationships were compiled across all nodes and converted to a NetworkX graph object, with optional restrictions on maximum nearest neighbors to simplify dense networks.

### Generate Nodes from Edge Vertices

Nodes were generated at branchpoints of binary edge segmentations using 3D skeletonization. Binary edge images were skeletonized using the sklearn.morphology.skeletonize() algorithm, with optional loop artifact correction through iterative hole filling and re-skeletonization. Terminal branches shorter than a specified pixel length were optionally removed by crawling from skeleton endpoints toward junctions. Branchpoints were identified by examining 3×3×3 neighborhoods around each skeleton voxel: the center voxel was set to zero and scipy.ndimage.label() was used to count distinct remaining elements, with locations having ≥3 distinct elements classified as branchpoints. Optional nearby branchpoints could be merged through dilation-based clustering. The resulting labeled branchpoint array was placed in the nodes channel for network analysis.

### Branchpoint Network Calculation

Branchpoint networks were calculated by first generating nodes from edge vertices (as described above), then establishing connections based on immediate spatial adjacency. Following branchpoint node generation, each node searched its immediate 3×3×3 neighborhood to identify which edges were encountered. Network connections were established between nodes that shared contact with the same edge segments, converting the branching structure into a node-based network representation suitable for graph analysis.

### Label Branches

Branch labeling was performed by combining branchpoint detection with skeleton segmentation. Following node generation from edge vertices, branchpoint nodes were used as masks to fragment the skeleton into discrete pieces. The fragmented skeleton pieces were labeled using scipy.ndimage.label() to assign unique identifiers to non-contiguous segments. Branch regions in the original binary image acquired labels from their corresponding skeleton pieces through distance transform-based assignment. Optional auto-correction steps included merging any internal labels with their external neighbors (from unusually thick regions having unpredictable skeletonization results), reassigned any branch labels that were non-contiguous in space to instead take on the label of their most-bordered neighbor, or reunified labels of branch segments that were of similar radii and trajectory in space (to optionally rejoin branch segments into more logical long branches).

### Branch Adjacency Network Calculation

Branch adjacency networks were calculated by combining branch labeling (as described above) with proximity network analysis. Following branch labeling to assign unique identifiers to discrete branch segments, a proximity network was calculated with a search distance of 1 voxel to identify immediately adjacent branches. This approach converted branching structures into networks where individual branch segments served as nodes, connected based on their spatial adjacency rather than through branchpoint connectivity.

### Centroid Calculation

Node and edge centroids were calculated as centers of mass for labeled objects. Images were optionally downsampled by a user-defined factor in all three dimensions to accelerate computation, with centroids subsequently normalized to full-resolution coordinates. Centroids for all labels were obtained via scipy’s center_of_mass() function.

## Image Processing Methods

**Resize:** Images were resized using scipy.ndimage.zoom with user-defined scaling factors applied to individual dimensions or all dimensions simultaneously. Cubic interpolation was optionally available for shape preservation but avoided for labeled data to maintain label integrity.

**Dilate:** Object dilation was performed using either pseudo-3D binary kernels or distance transform-based methods. Pseudo-3D kernels applied serial 2D dilations in XY and XZ planes using OpenCV2 algorithms to simulate 3D expansion, employed to enlarge objects for visualization but not for calculation reasons. Distance transform-based dilation inverted images, calculated scipy.ndimage.distance_transform_edt(), and applied boolean indexing for perfect dilations. Label-preserving dilation used distance transform indices to reassign dilated regions to their nearest source objects.

**Erode:** Object erosion employed similar algorithms to dilation but used OpenCV2 erode methods for pseudo-3D kernels and non-inverted distance transforms. Label-preserving erosion utilized skimage find_borders to identify object boundaries for proper label territory assignment during shrinkage.

**Fill Holes:** Holes were filled by iterating through 2D image planes (XY, and optionally YZ/XZ). In the case that borders were not designated for possible filling, the scipy.ndimage.binary_fill_holes method was used to fill the holes in each plane. If borders were considered, a slower method was used where each plane was inverted, then used scipy.ndimage.label() to identify contiguous background regions. Non-border regions were designated as holes and filled, with optional filling of small border holes sharing <8% border contact.

**Binarize:** Images were binarized either by setting all non-zero regions to 255 (total binarize) or using Otsu’s method for automatic foreground prediction and thresholding.

**Label Objects:** Connected component labeling assigned unique numerical identities to touching, non-zero regions using scipy.ndimage.label().

**Neighborhood Labels (or ‘Smart Label’):** Objects in binary images were labeled based on proximity to pre-labeled seed objects using distance transform indices to assign labels from the nearest labeled region. An alternate variant prioritized smooth labeling in binary regions contiguous with labeled ones, by first skeletonizing the binary image, then iterating through skeleton segments to assign these segments labels based on their overlap with the labeled image. This extended labeled skeleton was combined with the labeled image before the aforementioned distance-transform based method was applied. A final correction step could be used to reassign objects non-contiguous in space to have the labels of nearby objects they were actually contiguous with. The skimage.segmentation.watershed method could be deployed as an optional, slightly less accurate but often faster, way to flood fill regions by the labels of their neighbors.

**Threshold/Segment:** Intensity-based thresholding applied user-defined min/max value ranges to filter pixels by brightness values. Volume-based thresholding used pre-calculated object volumes for size-based selection. More generally, thresholding could be performed using any Python dictionary pairing numerical object labels to integer or float values (e.g., volumes, radii, network degree), allowing filtering based on any computed property that fits this label-value format.

**Machine Learning Segmentation:** Machine learning segmentation employed a Lightgbm classifier trained on user-designated positive and negative training regions. Two feature extraction modes were available, each utilizing scipy (or cupy, for GPU): “Quick Model” generated feature maps including the original image, Gaussian blurs (σ=1,2,4,8), difference of Gaussians (σ pairs: 1-2, 2-4, 4-8), and gradient magnitudes from scipy.ndimage.sobel kernels. “Detailed Model” additionally computed Gaussian gradient magnitudes for each σ, Laplacian of Gaussian second-order derivatives for each σ, and largest Hessian eigenvalues calculated from 3×3 Hessian matrices using second-order derivatives for each σ. Features were processed in 49³ voxel chunks for 3D neighborhoods or adaptively chunked 2D planes for images >64³ pixels. Intensity features remained in raw form while morphological features underwent z-score normalization. For color images, these feature maps were computed for each color channel and stacked together. Training regions provided labeled examples to the classifier, which then segmented entire volumes by applying the trained model to feature maps generated for each voxel location.

**Mask Channel:** One channel was used to create a binary mask applied to another channel, preserving only regions where the mask was non-zero.

**Crop Channels:** All channels were simultaneously cropped to user-defined spatial boundaries in X, Y, and Z dimensions.

**Channel dtype:** Image data types were converted between unsigned 8/16/32-bit integers and 32/64-bit floating point formats for memory optimization.

**Skeletonize:** 3D skeletonization used sklearn.morphology.skeletonize() to reduce objects to medial axes, with optional terminal branch removal and loop artifact correction through iterative hole filling and re-skeletonization. Terminal branches shorter than a specified pixel length were optionally removed by crawling from skeleton endpoints toward junctions. Since branch removal left holes at junction points, if this option was employed, the skeleton was dilated a single voxel to fill these holes, then re-skeletonized once more.

**Binary Watershed:** Watershed segmentation was used to split fused objects in binary images by identifying internal seed regions and growing them to object boundaries. A distance transform was computed on the binary image using scipy.ndimage.distance_transform_edt to calculate distances from each foreground pixel to the nearest background. Seed kernels were identified by thresholding the distance transform either by retaining only the top percentage of distance values (proportion method, e.g., top 5% for proportion=0.05) or by specifying a minimum radius threshold to exclude objects smaller than the specified size. The proportion method excluded distance values below the specified percentile, while the radius method provided more intuitive control by directly specifying the smallest object radius to retain. These seed regions were then labeled using scipy.ndimage.label() to assign unique identifiers to each disconnected seed. Finally, the skimage.segmentation.watershed algorithm grew these labeled seeds outward to segment the original binary objects, effectively separating touching or overlapping structures into distinct labeled components.

**Gray Watershed:** Grayscale watershed segmentation separated objects in intensity images by identifying local intensity maxima and using them as growth seeds. The skimage.feature.peak_local_max function identified intensity peaks within the foreground-segmented image, with peaks required to be separated by a user-defined minimum distance to prevent over-segmentation of single objects. An additional minimum peak intensity threshold could be applied to ignore low-intensity regions during peak detection. The identified peaks were marked on a copy of the image array to serve as labeled seed points. The original grayscale image was then processed using the skimage.segmentation.watershed algorithm, which grew the seed regions outward following intensity gradients until they met at watershed boundaries, effectively segmenting connected objects based on their intensity profiles and the spatial distribution of local maxima.

**Invert:** Images were inverted by reversing intensity values (high to low, low to high).

**Z-Project:** 3D image stacks were collapsed into 2D projections using maximum, mean, minimum, sum, or standard deviation operations across the Z-dimension.

**Normalize Brightness:** To correct illumination gradients in three-dimensional microscopy volumes, we apply a sequential, shell-based normalization along each spatial axis. For a given axis, a two-dimensional Euclidean distance transform (EDT) is computed on each orthogonal tissue-mask slice, and the resulting depth values are partitioned into *n* equal-width concentric shells. Within each shell, a one-dimensional median-intensity profile is computed along the target axis by collapsing across the two orthogonal axes, and per-position multiplicative correction factors are derived as the ratio of the shell’s global median to the local profile value. Corrections are applied sequentially for each axis (Z, Y, X).

**Generate Nodes from Node Centroids:** Node images were generated from pre-existing centroid coordinates stored in the node_centroids property. Each centroid was converted to a labeled point in a new image array, with image dimensions bounded by the maximum centroid coordinates in each dimension. The resulting labeled point image was placed in the nodes channel, enabling access to image-based functions when only centroid data was available from previous sessions or external analysis tools.

**Trace Filaments:** Tiny connected components (< 10 voxels) were first filtered of a binary segmentation. The remaining segmentation had its distance transform obtained via scipy.ndimage. distance_transform_edt, to evaluate the distance of the background to its skeleton as a measurement of radii. The remaining segmentation was likewise skeletonized, followed by hole-filling followed by another round of skeletonization to account for looping artifacts in 3D images. A series of kernel points were then placed along the skeleton a user-specified distance apart, including all endpoints. For each kernel, geometric features such as radius and direction were obtained. By crawling along the skeleton, all internal kernels were connected to their immediate neighbors to build a NetworkX graph. Any endpoints evaluated nearby kernels for potential connections, scoring them based on similar kernel characteristics, such as radii, trajectory, and surrounding features. Those with a high enough score were connected in the NetworkX graph. Next, connected components in the graph that showed geometric likelihood of being noise (such as having a blob-like structure) were removed. Finally, each kernel had a tapered cylinder drawn to its neighbors, based on their calculated radii, resulting in a smoothed segmentation. As an optional final step, components of the resulting segmentation that were both above a specified sphericity and a specified volume could be filtered out, as such blobs were likely noise.

**Generate Voronoi Diagram:** Voronoi diagrams were generated from node centroids using a distance transform-based approach equivalent to smart dilation with maximal search distance. Each image voxel was assigned the label of its nearest centroid, creating tessellated regions where each cell represented the spatial territory closest to a given node. The resulting Voronoi diagram was loaded into Overlay2 and could serve as an alternative method for defining node neighborhoods in connectivity network calculations, particularly suitable for small or homogeneous spheroidal nodes.

### Generate Convex Hull

The function extracted coordinates of all foreground voxels via np.argwhere, computed their convex hull, then built a Delaunay triangulation over the hull vertices only. A full coordinate grid spanning the array was constructed with np.indices, and Delaunay.find_simplex classified every grid point simultaneously. These were written into an output mask.

### Generate Hexagons/Rhomboids

Seed points were placed at the theoretical centers of a hexagonal lattice in 2D, or at Face-Centered Cubic lattice positions in 3D — the FCC lattice being the natural generator of rhombic dodecahedral Voronoi cells. Each seed was a single marked voxel in an otherwise empty array, then scipy.ndimage.label assigned each seed a unique integer. distance_transform_edt was then called on the complement (unlabeled voxels), returning not distances but the coordinates of the nearest labeled voxel for every point in the volume; indexing the seed-label array with those coordinates propagates region membership to all voxels — equivalent to computing a Voronoi diagram but without any explicit nearest-neighbor search. The resulting tessellation was hexagonal in 2D and rhombic-dodecahedral in 3D purely by virtue of the lattice geometry, since Voronoi cells of a hexagonal and FCC lattice were exactly those shapes respectively. The resultant hexagons/dodecahedrons could be confined to a mask region to serve as artificial nodes for neighborhood characterization.

## Modify Network/Properties Methods

**Remove Unassigned IDs from Centroid List:** Centroids lacking associated identity assignments were removed from the node_centroids property to ensure compatibility with ID-dependent functions.

**Force Multiple IDs to Single ID:** Nodes with multiple identity assignments were randomly assigned a single identity to simplify identity visualization and reduce permutation complexity.

**Remove Nodes Not in Nodes Channel:** Node entries were removed from node_centroids and node_identities properties if their corresponding labels were absent from the current nodes channel image, primarily used after image cropping operations.

**Remove Trunk:** The most highly interconnected edge (trunk) was identified and removed from the network to eliminate dominant central structures and reveal downstream connectivity patterns.

**Convert Trunk to Node:** The trunk edge was converted to a new node rather than removed, preserving network structure by treating the trunk as a central hub while transferring the trunk region from the edges image to the nodes image.

**Convert Edges to Node Objects:** All edges were converted to nodes, merging the edge and nodes images while updating the network to connect nodes to their previously shared edges. Edge objects received new non-overlapping labels and were assigned “edge” identities in the node_identities property.

**Remove Network Weights:** Edge weights representing multiple connections between node pairs were removed, reducing the network to binary connectivity relationships.

**Prune Same-Type Connections:** Connections between nodes sharing identical identities were removed, retaining only inter-type relationships when node_identities were defined.

**Isolate Specific Type Connections:** Networks were filtered to retain only connections involving two user-selected node identity types, removing all other node relationships.

**Rearrange Community IDs by Size:** Community identifiers were reassigned in descending order of community size, with the largest community receiving ID=1 and subsequent communities numbered sequentially.

**Convert Communities to Nodes:** Node-level networks were converted to community-level networks, with individual nodes in the nodes channel taking on their community labels rather than original identifiers.

**Add/Remove Network Pairs:** Manual network editing allowed addition or removal of specific node pairs, with optional edge ID specification for precise connection management.

## Assignment of Node Identities and Related Analysis

### Node Merging Algorithm

To enable comparison of heterogeneous node populations (e.g., different cell types or structural elements), individually segmented and labeled node images were merged into a single composite channel. Prior to merging, centroids were optionally computed for each input image to preserve accurate spatial coordinates regardless of potential label conflicts in the merged output. An optional downsampling factor could be applied during centroid calculation to improve computational efficiency, though this risked loss of small nodes. Node identities were automatically assigned based on the source image filename, with the original nodes designated as ‘root_nodes’. The merged composite retained non-overlapping labels from all input images, enabling simultaneous analysis of multiple node populations within the same network framework.

### Node Identity Assignment from Image Overlap

Node identities were assigned based on spatial overlap with additional imaging channels. Nodes could search outward from their centroid positions by a user-defined step-out distance to account for cell radius or morphological extent, computed via distance transform. Spatial scaling parameters (xy_scale and z_scale) converted pixel distances to biological units when specified. Two binarization strategies were available: automatic thresholding using either Otsu’s method to distinguish foreground from background or using presegmented binary channels, or manual thresholding based on user-guided mean intensity overlap between nodes and each channel. For manual thresholding, optional UMAP visualization displayed intensity Z-scores relative to channel identity. Nodes exceeding the threshold for a given channel received a positive identity marker (+), while those below threshold could optionally receive negative markers (-). Multiple channel identities were concatenated into composite identity strings for each node, enabling phenotypic classification of heterogeneous node populations.

### Thresholding via Neural Network

While doing manual threshold assignment of the above, the mean intensity of all cells in the image was used to generate a 70-bucket histogram, where the first 49 buckets were smaller to capture the majority of relevant data, confining any long tail to the last 21 buckets. These histograms were used to train a neural network to associate histogram features with desired min/max threshold bounds. Raw histograms were converted to slope arrays (finite differences between adjacent bins normalized by bin width), capturing the shape of the intensity distribution more naturally than raw counts. These slopes were concatenated with their corresponding bin boundary positions to form a fixed-length feature vector per channel, which is then standardized via standardscaler. An MLP regressor (PyTorch or sklearn backend, auto-selected by hardware availability) was trained to map these feature vectors to [min, max] threshold pairs, with both inputs and outputs independently z-scored so the network learned in normalized space before predictions were inverse-transformed back to raw intensity values. The model supported incremental retraining via warm-start, and serialization stored weights in both PyTorch and sklearn-compatible formats simultaneously, enabling cross-backend loading with automatic weight transposition to reconcile the differing [in, out] vs [out, in] storage conventions between the two frameworks.

### Violin Plots of Channel Intensities for Node Identities

To characterize the expression profiles of identified node populations, normalized violin plots were generated from average intensity values for each node across all marker channels. These values were extracted from the identity assignment workflow and subjected to a modified Z-score normalization procedure. For each channel, normalization was centered around the minimum intensity value observed among nodes positively identified for that marker, rather than the population mean. This approach quantified the degree to which nodes deviated from the baseline threshold defining positive marker expression. For channels without corresponding node identities, the population median served as the normalization reference point. Each violin plot represented the expression profile of a single channel across all nodes within a specified identity or neighborhood. Values greater than zero indicated expression levels above the validated threshold for that marker, enabling assessment of co-expression patterns across multiple channels.

### UMAP Visualization Channel Intensities of Node Identities

Dimensionality reduction for visualization was performed using Uniform Manifold Approximation and Projection (UMAP) on standard Z-score normalized intensity data. Prior to UMAP generation, each node’s intensity values across all channels were normalized using sklearn’s StandardScaler, centering values around the population mean and scaling by standard deviation. Nodes in the resulting UMAP space were labeled according to either their assigned identities or their computed neighborhood memberships, facilitating visual assessment of population separation and phenotypic heterogeneity.

### Neighborhood Assignment based on Channel Intensities for Nodes

For unsupervised neighborhood assignment, nodes were clustered based on their multi-channel intensity signatures using K-means clustering (sklearn’s KMeans). Intensity values were normalized using standard Z-score transformation prior to clustering. The optimal number of clusters was either user-defined or computationally predicted via Calinski-Harabasz scoring (maximum k=20). Each resulting neighborhood was characterized by its aggregate intensity profile across all channels.

## Network Visualization and Community Partitioning

### Network Visualization

Networks were visualized using custom pyqtgraph interface with several layout options: centroid layout positioning nodes according to their 3D spatial coordinates (with XY position corresponding to image location and node size representing Z-position), spring layout for force-directed positioning, shell layout for visualization of connectivity to a most-central node, with optional separation of graph components. Visualization modes included default numerical node IDs, community-based color coding, or node identity-based color coding. Generated graphs could be automatically saved to specified output directories.

### Generic Network Report

Basic network statistics were computed including node count, edge count, density, connectivity status, connected components, degree distributions (average, maximum, minimum), centrality measures (betweenness, closeness, eigenvector), clustering coefficient, transitivity, diameter, average shortest path length, triangle count, degree assortativity, and identification of unconnected nodes. Additional statistics included node counts per identity category and community membership when available. All statistics were generated using NetworkX functions and displayed in tabulated format.

### Network Community Detection and Statistics

Network communities were identified using either label propagation or Louvain algorithms, with optional consideration of edge weights. Community statistics included network-level metrics (modularity, community count and sizes, global clustering coefficient, assortativity, inter-community edges, mixing parameter) and community-specific metrics (density, conductance, average clustering, degree centrality, average shortest path length). Random seed parameters enabled reproducible partitioning. Detected communities were saved as node properties and exported in tabulated format alongside statistical summaries. An alternative, non-network approach split the image into approximate cubes of user-defined side length and assigned communities based on which node fell into which cube. This option was available for when spatial analysis was still desired but nodes were arranged randomly, which would result in meaningless networks.

### Community-to-Supercommunity Conversion

Network communities were consolidated into larger spatial supercommunities through a multi-step clustering process that analyzed community identity compositions and grouped similar communities together. The method began by calculating the proportional makeup of different node identity types within each community using the identity composition analysis algorithm. These compositional profiles were then converted into numpy arrays suitable for scikit-learn clustering algorithms.

Two clustering approaches were available: K-means clustering (generally recommended) and DBSCAN clustering (experimental). For K-means clustering, users could specify the desired number of supercommunities or allow automatic determination through Calinski-Harabasz scoring evaluation of supercommunity counts from 1 to 20, selecting the configuration with the highest quality score. Random seeds (default: 42) ensured reproducible clustering results. DBSCAN clustering autonomously determined supercommunitiy numbers using calculated parameters: minimum samples set to max(3, √n_samples × 0.2) for core point identification, and epsilon values derived from the 80th percentile of 4th nearest supercommunity distances to define supercommunity radius. Points failing to meet core or border criteria became outliers assigned to “Supercommunity 0.”

### Neighborhood Community Detection

For each node in the graph, its immediate neighbors were retrieved and their cell-type identities accumulated into a fixed-length count vector (one dimension per identity in the global set), then normalized by neighbor count to yield a composition fraction profile — making each node’s feature vector scale-invariant with respect to degree. Nodes with fewer neighbors than a user-defined minimum threshold were excluded. These neighborhood composition vectors are then clustered via KMeans into a user-specified number of communities, grouping nodes whose local cellular microenvironments are compositionally similar regardless of spatial position. Cluster assignments are unpacked and re-keyed by node, then optionally characterized by averaging the composition vectors of all member nodes to produce per-cluster heatmaps reflecting the mean neighborhood phenotype of each community.

### Channel Intensity Community Detection

Utilized a similar set of steps as ‘Neighborhood Community Detection’, however vectors were computed based on normalized intensity (via StandardScaler) per marker per node. The resultant feature vectors between all nodes were clustered via KMeans into a user-specified number of communities.

### Community Identity Composition Analysis and UMAP Visualization

Community compositions were characterized by analyzing the proportional makeup of node identity types within each detected community. Two analysis modes were available: individual community analysis providing compositional profiles for each community separately, or weighted average analysis computing network-wide identity distributions weighted by community size to emphasize larger communities. For individual community analysis, the method calculated the proportion of each node identity type within every community and optionally generated UMAP visualizations to reveal compositional similarities between communities.

UMAP generation utilized the Python umap module with fixed random seed (42) for reproducible dimensionality reduction, transforming high-dimensional community composition vectors into 2D coordinates for visualization. Communities were positioned in UMAP space such that those with similar identity compositions clustered together, enabling identification of functionally similar network regions. UMAP points could be unlabeled, labeled by community number, or colored by neighborhood assignment when communities had been previously converted to neighborhoods.

Two compositional analysis approaches were available through the “Return Node Type Distribution Robust UMAP” option. Standard analysis grouped communities by their raw proportional identity compositions. Robust analysis instead evaluated communities based on identity overrepresentation, where communities were characterized by how much they overrepresented specific node types compared to random distribution expectations. Overrepresentation was calculated as the ratio of observed identity proportion within the community to the expected proportion based on overall network composition. Communities below a minimum size threshold could be excluded from UMAP analysis to focus on statistically significant clusters.

For the alternative output of this method (weighted average analysis – which describes a prototypical ‘average community’), communities were grouped by identity type, identity counts were weighted by community size, summed across all communities, and normalized first by total node count then to ensure proportional summation to unity. This approach provided network-level compositional metrics that emphasized the contribution of larger, more significant communities while maintaining sensitivity to overall network heterogeneity.

## Network Statistical Analysis

### Generic Network Statistics Calculation

Comprehensive network statistics were computed using NetworkX functions to characterize overall network topology and connectivity patterns. Basic structural metrics included node count, edge count, network density, directedness status, connectivity status, number of connected components, and largest component size.

Degree-based measures encompassed average, maximum, and minimum node degrees alongside degree assortativity coefficients. Centrality analyses calculated average betweenness centrality (identifying bridge nodes), closeness centrality (measuring reach efficiency), and eigenvector centrality (assessing prestige through high-quality connections). Clustering properties were evaluated through average clustering coefficients and transitivity measures. Path-based statistics included network diameter and average shortest path length between all node pairs. Additional topological features assessed included tree structure identification, triangle enumeration, and detection of unconnected nodes excluded from the original node image. All computed statistics were exported to tabulated data widgets for further analysis and comparison across different network configurations.

### Network Statistics Distribution Analysis

Network node properties were characterized through histogram distributions of various centrality and connectivity measures computed using NetworkX algorithms. Available distributions included degree distribution (connection count per node for identifying hub-based versus egalitarian network architectures), shortest path length distribution (minimum steps between node pairs for assessing network efficiency and small-world properties), and multiple centrality measures. Centrality distributions encompassed degree centrality (direct influence through immediate connections), betweenness centrality (identification of critical bridge nodes), closeness centrality (optimal positioning for information spreading), eigenvector centrality (prestige through connections to important hubs), harmonic centrality (robust closeness measure for disconnected components), load centrality (traffic burden assessment), current flow betweenness (electrical circuit-based flow modeling), and communicability betweenness (multi-step communication importance).

Additional distributions analyzed clustering coefficients (neighbor interconnectedness), triangle counts (local group cohesion), k-core decomposition (hierarchical dense subgroup identification), eccentricity (maximum distances to other nodes), node connectivity (local robustness assessment), average dispersion (neighbor scatter measurement), and network bridge identification (critical connection detection). All distributions were visualized as matplotlib histograms with corresponding numerical data exported to tabulated formats for quantitative analysis.

### Radial Distribution Analysis

Network connectivity patterns were characterized as a function of spatial distance through radial distribution analysis, which quantified the average number of neighboring nodes at incrementally increasing distances from any given node position. The method systematically searched outward from each node in the network using user-defined distance buckets as step sizes, counting connected neighbors within each radial shell. Distance measurements utilized the 3D spatial coordinates of node centroids, with bucket distances scaled according to image calibration parameters when specified. The analysis generated distance-versus-neighbor count profiles that revealed spatial organization patterns of network connectivity, enabling assessment of connection efficiency and identification of preferential connection distances. Networks with efficient spatial organization typically exhibited abundant short-range connections with fewer long-range connections, producing characteristic decay profiles in the radial distribution curves. Results were visualized as matplotlib line graphs plotting average neighbor count against radial distance, with corresponding numerical data exported to tabulated formats. Output graphs could be automatically saved to specified directories for documentation and comparative analysis across different network configurations.

### Community Cluster Heatmap

Community spatial density patterns were visualized through heatmap analysis that compared observed community sizes to expected random distribution densities across the network. The method calculated baseline density expectations by dividing the total number of nodes by the number of communities, representing the expected community size under random uniform distribution. Heat values for individual communities were computed using natural logarithm ratios of actual community size to expected baseline size, generating normalized density metrics that identified communities significantly larger or smaller than random expectations.

Communities with densities above baseline expectations were represented in red coloration, indicating unexpectedly dense clustering, while communities below expectations appeared in blue, signifying sparse or dispersed organization. The logarithmic transformation ensured symmetric representation of over– and under-density patterns while maintaining sensitivity to moderate deviations from random expectations.

Visualization options included 2D or 3D matplotlib scatter plots positioned according to node centroid coordinates, with point colors mapped to calculated heat values using standardized color scales. Alternative overlay generation created RGB image overlays deposited into Overlay2 channel for integration with other imaging data and spatial context preservation. Total node counts could be manually specified to accommodate filtered datasets where active nodes represented subsets of complete populations, ensuring accurate density baseline calculations.

The method required both node centroids and community assignments, automatically prompting for property generation when unavailable. Output included both visual heatmaps and tabulated data showing community identifiers paired with corresponding density intensity values, enabling quantitative assessment of community organization patterns and identification of spatially anomalous clustering behaviors across the network structure.

## Non-Network Spatial Analysis

### Identity Distribution of Neighbors

Neighborhood composition analysis characterized the spatial relationships between different node identity types through two distinct approaches: network-based connectivity analysis and morphological proximity analysis. Network-based analysis (Mode 1) examined immediate network neighbors connected through graph edges, quantifying both absolute neighbor counts and proportional representation of each identity type relative to the total available nodes of that type across the network. This approach revealed functional connectivity patterns between different node populations based on actual network topology.

Morphological proximity analysis (Mode 2) evaluated spatial neighborhoods within user-defined search radii around nodes of a specified root identity type, excluding the root nodes themselves from analysis. Distance-based searching utilized precise distance transforms for neighborhood identification. This mode generated three complementary metrics: total volumes of each identity type within the specified radius (absolute spatial proximity), proportional representation of each identity type’s total network volume within the search radius (relative spatial clustering), and clustering factors comparing observed identity densities to expected random distribution densities.

Clustering factors served as relative density measures, with values greater than 1 indicating preferential spatial association (overrepresentation) between the root identity and target identity types, while values less than 1 suggested spatial avoidance (underrepresentation) compared to random distribution expectations. Both analysis modes generated corresponding bar chart visualizations alongside tabulated data exports, enabling quantitative assessment of identity-specific neighborhood compositions and identification of preferential spatial or functional associations between different node populations within the network structure.

### Ripley’s K Clustering Analysis

Spatial clustering patterns were quantified using Ripley’s K function analysis, which compared observed object distributions to theoretical Poisson (random) distributions across multiple distance scales. The analysis evaluated clustering behavior either between different node identity types or within single identity populations by constructing KDTree data structures for efficient nearest-neighbor searching. Root points (nodes of specified identity) were systematically queried for target points (same or different identity) within incrementally increasing radii defined by user-specified bucket distances, with spatial scaling parameters converting pixel distances to biological units when provided.

For each distance radius, the method calculated K values by summing neighbor counts across all root points, normalizing by point density (λ = number of points / study region volume) and root point count. K values were subsequently transformed to H values using dimensional-specific formulas: H = √(K/π) – r for 2D data or H = ∛(K/(4π/3)) – r for 3D data. Results were plotted against theoretical expectations (H = 0 for random distributions), with positive deviations indicating spatial clustering and negative deviations suggesting dispersion or regularity.

Multiple border correction strategies addressed edge artifacts inherent in finite study regions. Internal node restriction limited analysis to nodes beyond specified distances from image boundaries or within user-defined tissue masks created from binary foreground channels (with distance measurements derived from its distance transform. Point reflection correction extended the analysis region by mirroring points across image boundaries, creating up to 8 mirror regions (2D) or 26 mirror regions (3D) to simulate extended spatial domains. Distance transform-based masking enabled restriction of analysis to biologically relevant tissue areas while maintaining accurate density calculations.

Additional parameters controlled analysis robustness, including proportional image search limits (0-1 range) to balance accuracy against border artifacts, and boundary definition options using custom tissue masks for irregular specimen geometries. Results generated both tabulated K and H values alongside matplotlib visualizations comparing observed clustering patterns to theoretical random distributions, enabling quantitative assessment of spatial organization across multiple distance scales.

### Average Nearest Neighbor Analysis

Nearest neighbor relationships were quantified through distance measurements between nodes, with analysis options for identity-specific interactions when node identity properties were assigned. The method supported two distance calculation approaches: centroid-based analysis using point coordinates for rapid processing of spherical objects, or object boundary-based analysis using complete object perimeters detected via skimage boundary identification for complex morphologies.

Users specified the number of nearest neighbors for averaging (default: 1), with higher values providing smoothed distance estimates by averaging multiple proximal relationships per node. However, in the case that object boundaries were used, the number of nearest neighbors was limited to 1 to avoid computational strain. Furthermore, instead of just querying between distinct points, the object boundary method first computed the coordinates of the image boundaries via skimage.segmentation.find_boundaries method. The boundary points of each labeled object in the root nodes were used to search for the nearest point of the target nodes.

Distance calculations utilized KDTree data structures built from neighbor node coordinates for efficient spatial querying. For each root node, the specified number of nearest neighbors were identified and their distances averaged to generate per-node proximity metrics. When node identities were assigned, analysis could focus on specific identity type interactions, examining distances from root identity nodes to target identity populations, including self-interactions or relationships excluding the root type.

Heatmap generation provided spatial visualization of neighbor distance patterns, with color intensity based on comparison to theoretical uniform distribution expectations. The expected uniform distance was calculated by distributing target points evenly throughout the image space (or within user-defined tissue masks) and measuring distances using identical neighbor-finding parameters (by re-simulating the KDTree centroid calculation, in the uniform space). Heat values utilized logarithmic ratios: ln(expected uniform distance / observed distance), with red coloration indicating closer-than-expected proximity and blue indicating greater-than-expected separation.

Output options included tabulated distance distributions for individual nodes, averaged distance summaries across all identity combinations, 2D or 3D matplotlib heatmaps, quantifiable grayscale overlays with pixel intensities equal to measured distances (Overlay1), and RGB color-coded heatmaps (Overlay2). All distance measurements incorporated xy_scale and z_scale calibration parameters to ensure biologically relevant distance units, with automatic prompting for property verification when spatial scaling was critical for analysis accuracy.

### Node-Edge Interaction Analysis

Spatial relationships between node and edge channels were quantified by measuring edge channel intensities within defined search distances from each labeled node object. The analysis employed the same computational framework as radii calculation, using find_objects() for bounding box identification and subarray extraction from both node and edge channels, followed by distance transforms via distance_transform_edt() for search region measurements.

Search regions could include or exclude internal node areas depending on analysis objectives. For exclusion mode, inverted boolean arrays removed core node regions from dilated search masks, isolating edge interactions to node peripheries. Edge intensities within search regions were summed and volumetrically scaled using dimensional calibration parameters. The parallel processing framework ensured efficient computation across large datasets while maintaining per-object specificity for detailed interaction characterization. Results were returned as simple volumes, or as lengths, which were calculated by skeletonizing the surrounding edges, followed by applying the distance formula between each adjacent skeleton voxel.

## Morphological Analysis

**Volume Calculation** Object volumes were quantified using numpy bincount operations applied to labeled image arrays, with voxel counts scaled by axis calibration parameters (xy_scale, z_scale) to convert pixel measurements to biological volume units. The method processed all labeled objects in the active image channel simultaneously, generating comprehensive volume datasets exported to tabulated formats.

**Radii Calculation** Maximum object radii were determined through distance transform analysis using scipy.ndimage methods. Object bounding boxes were identified via find_objects(), with padded subarrays extracted around each labeled region. Within each subarray, boolean indexing isolated individual objects, followed by distance_transform_edt() application to generate distance maps. Maximum distance values represented the largest radii (furthest points from object boundaries). The process utilized parallel processing across all available CPU cores for computational efficiency

**Approximate Surface Area Calculation** Surface areas of segmented objects were calculated using a voxel-based boundary detection method. The algorithm treated each voxel as a discrete cube and identified exposed faces by counting the number of adjacent occupied voxels in the 3D grid. External faces (those not adjacent to another voxel of the same object) were enumerated using numpy’s roll and bincount functions to efficiently scan across all three spatial dimensions. This discrete voxel-based approach resulted in a tessellated surface representation that overestimates surface area for objects with smooth boundaries, as all surfaces were approximated as stepped rather than continuous. This method was selected to prioritize computational efficiency for rapid analysis of large datasets.

**Sphericity Calculation** To quantify the geometric regularity of segmented objects, sphericity values were calculated for all objects in each image. Sphericity (ψ) is a dimensionless shape descriptor ranging from 0 to 1, where a perfect sphere yields ψ = 1 and increasingly irregular shapes approach 0. Sphericity was computed using the standard formula:

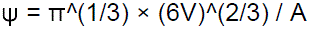

where V is the object volume and A is the surface area. This metric provides a quantitative assessment of how closely each object approximates an ideal sphere, independent of object size.

**Branch Tortuosity And Length Calculation** Branch length was calculated by summing the Euclidean distances between adjacent skeleton voxels for each labeled branch. Tortuosity was quantified as the ratio of actual branch length to the maximum straight-line distance between branch endpoints.

## Data and Overlay Generation Methods

**Degree Information Analysis** Node connectivity patterns were visualized and quantified through degree-based overlay generation and tabulation. Four execution modes were available: tabulated degree data without visualization, degree value text overlays drawn at node centroids, node relabeling by degree values for downstream thresholding, and RGB heatmaps with red indicating above-average connectivity and blue representing below-average connectivity. Optional filtering retained only high-degree nodes based on user-specified proportional thresholds (0-1 range), enabling focus on network hubs. Temporary downsampling accelerated overlay creation for large datasets while maintaining analytical accuracy.

**Hub Node Identification** Network hub nodes were identified as those with minimal degrees of separation to all other nodes within their connected components, representing the most centrally positioned nodes for information flow. Hub analysis operated independently within each network component to avoid bias from component size variations. Proportional thresholds (0-1 range) enabled selection of top-ranking hubs, with components containing insufficient nodes excluded when thresholds were restrictive. Optional overlay generation highlighted identified hub nodes in the Overlay2 channel for spatial visualization of network control points.

**Mother Node Detection** Inter-community bridge nodes (“mother nodes”) were identified as nodes maintaining connections spanning multiple communities, enabling communication between otherwise separated network regions. The analysis required pre-existing community assignments and systematically evaluated each node’s connectivity pattern across community boundaries. Mother nodes were tabulated with their inter-community connection profiles and optionally highlighted in Overlay1 channel overlays for spatial identification of critical network bridges facilitating cross-community interactions.

**Community and Identity Coding** Spatial visualization of community and identity assignments utilized color-coded or grayscale overlay generation. Color-coded mode created RGB overlays with distinct colors assigned to each community or identity type, providing intuitive visual discrimination of node populations. Grayscale mode assigned numerical labels corresponding to community or identity classifications, enabling downstream thresholding and quantitative analysis of specific populations. Legend tables accompanied all coding overlays, mapping numerical or color assignments to their corresponding classifications. Downsampling options improved computational efficiency for large datasets.

**Centroid UMAP Visualization** Spatial organization patterns were revealed through UMAP clustering based on node centroid coordinates, identifying nodes with similar spatial positioning within the 3D image volume. The method was particularly valuable for 3D datasets where spatial relationships were difficult to assess through direct visualization. When node identities were assigned, points were colored according to identity classifications, with unassigned nodes labeled as “Unknown.” The resulting 2D embedding revealed spatial clustering patterns and neighborhood relationships that reflected underlying tissue organization or experimental spatial structure.

## Randomization Methods

**Equivalent Random Network Generation** Control networks were generated with identical node and edge counts to the original network while randomizing connectivity patterns to establish null distribution baselines for comparative analysis. The method preserved total edge counts, including weighted edge contributions when present, but redistributed connections randomly among available nodes. For weighted networks, users could specify whether the random network could utilize weighted edges, with edge weights consuming proportional connection allowances from the total edge budget. This approach enabled assessment of non-random connectivity patterns by comparing observed network properties against equivalent random networks with identical structural constraints but randomized topology.

**Node Spatial Randomization** Node spatial distributions were randomized while preserving node counts and identity compositions to evaluate spatial organization patterns independent of network connectivity. The method utilized node centroids for position randomization, with five spatial constraint options: unrestricted placement anywhere within image boundaries, placement within the existing node bounding box, restriction to non-zero edge channel regions, or confinement within non-zero Overlay1 or Overlay2 channel areas. Custom boundary options enabled creation of biologically relevant spatial constraints by dilating regions of interest to define permissible node locations. When node channel images existed, they were replaced with equivalently-sized randomized distributions. For datasets lacking node images, only centroid coordinates were randomized within available image boundaries or existing centroid ranges. This spatial randomization enabled assessment of clustering patterns, neighborhood compositions, and distance-based relationships against spatially random null distributions while maintaining node population characteristics.

## Significance Testing GUI

Statistical comparisons from the NetTracer3D GUI were performed using the scipy.stats module in Python. Available tests included: Student’s t-test for paired and independent samples (stats.ttest_rel and stats.ttest_ind with equal_var=True, respectively), Welch’s t-test for independent samples with unequal variances (stats.ttest_ind with equal_var=False), one-way ANOVA for comparing multiple groups (stats.f_oneway), and the non-parametric Mann-Whitney U test (stats.mannwhitneyu). Correlation analysis was conducted using Pearson’s correlation coefficient (stats.pearsonr). Normality of data distributions was assessed using the Shapiro-Wilk test (stats.shapiro). For categorical data, the chi-square test of independence was performed using stats.chi2_contingency with contingency tables generated via pandas crosstab. Effect sizes for chi-square tests were calculated using Cramér’s V. Statistical significance was determined at α = 0.05 for all tests.

## Image Menu Methods

### Brightness/Contrast Adjustment

Image visualization was optimized through an adaptive brightness/contrast adjustment algorithm that operated on 16-bit grayscale data (0-65535 range). Each channel was independently adjustable using dual-threshold sliders, allowing users to define minimum and maximum brightness values for optimal contrast enhancement. This preprocessing step was essential for visualizing low-contrast biological structures and ensuring consistent image quality across different acquisition conditions.

### Channel Color Assignment

The software implemented a flexible color mapping system where each data channel could be assigned distinct colors from a predefined palette. Default assignments included light red for node channels, light green for edge channels, and white/cyan for overlay channels. This color-coding system enhanced visual discrimination of different structural components during analysis and visualization.

### Network Overlay Generation

Spatial network connectivity was visualized through an automated overlay generation algorithm that created 1-voxel thick white lines connecting all node centroids. The method required pre-computed node centroid coordinates and generated vector-based network representations placed in Overlay1. Line thickness could be enhanced through morphological dilation operations for improved visibility in large-scale datasets.

### Node Identification Overlay

Individual node identification was facilitated through an automated labeling system that overlaid numerical identifiers directly onto node centroid positions. This method generated text-based overlays in Overlay2, providing immediate visual correspondence between network analysis results and spatial locations within the image data.

### Object Colorization Algorithm

The software employed a unique color assignment algorithm that converted grayscale-labeled objects (nodes or edges) into distinct RGB color representations. Each grayscale intensity value was mapped to a unique color from an internal palette, generating comprehensive color-coded overlays in Overlay2. A corresponding legend was automatically generated, documenting the color-to-object correspondence. An optional downsampling parameter allowed processing optimization while maintaining object integrity.

### Object Selection and Navigation

The software provides targeted object selection capabilities through a search algorithm that accepts comma-separated integer lists corresponding to object identifiers within either the nodes or edges channels. Selected objects are visually highlighted within the image display, and the system automatically navigates to the z-plane containing the centroid of the first selected object, enabling rapid spatial localization of specific network components. The selection system supports both manual entry of object identifiers and automated import from external spreadsheet files (.csv or .xlsx format), where object identifiers are extracted from the first column. A complementary deselection function allows removal of objects from the current selection set, with selection operations taking precedence over deselection when the same object appears in both lists. This functionality supports complex analysis workflows including quality control procedures, subset analysis, and systematic examination of network components organized through external classification systems.

### Three-Dimensional Visualization

NetTracer3D integrated with the Napari^17^ visualization platform to provide GPU-accelerated 3D rendering capabilities. The method supported multi-channel visualization with optional downsampling for performance optimization. Cubic resampling algorithms could be used to preserve object morphology during downsampling operations. Optional bounding box rendering provided spatial reference within the 3D environment.

### Cellpose3 Integration

The software provided direct integration with the Cellpose3 segmentation platform^11^, automatically launching the appropriate interface based on image dimensionality (2D or 3D). This integration enabled seamless transition between NetTracer3D’s network analysis capabilities and state-of-the-art deep learning-based cellular segmentation, supporting comprehensive image analysis workflows.

## NetTracer3D Right-Click Interactive Methods

### Image Viewer Window Right-Click Functions

#### Object Search and Navigation

The software implemented a direct object search algorithm accessible through right-click context menus. Users could specify target object identifiers (nodes, edges, or communities) through a text input interface, with automatic selection, highlighting, and z-plane navigation to the object’s calculated centroid position. A dropdown menu allowed selection between different object types for targeted searching.

#### Network Neighbor Analysis

Neighborhood analysis functions identified and visualized first-degree network connections through graph traversal algorithms. Three operational modes were supported: node-only neighbor selection, combined node-edge neighbor selection, and edge-only neighbor selection. The algorithms traversed the network topology to identify all objects within one degree of separation from currently selected elements, adding them to the active highlight display.

#### Connected Component Identification

Connected component analysis employed graph connectivity algorithms to identify and select all elements within the same network subgraph. The method supported three visualization modes: nodes only, combined nodes and edges, or edges only. The algorithm performed breadth-first traversal from selected objects to identify all reachable network elements within the same connected component.

#### Community Visualization

Community-based selection utilized pre-computed community partition data to enable group-wise object selection. The method supported both node-only and combined node-edge selection modes, identifying all network elements assigned to the same community as currently selected objects. This functionality required prior execution of community detection algorithms.

#### Identity-Based Selection

Identity classification enabled categorical object selection based on user-defined node identity assignments. The system provided dynamic menu options corresponding to each defined identity class, allowing bulk selection of all nodes sharing the same identity classification.

#### Comprehensive Selection Tools

Bulk selection algorithms provided complete object selection capabilities across three modes: all nodes, all edges, or combined selection of both object types. These methods enabled rapid selection of entire object populations for subsequent analysis operations.

#### Object Manipulation Operations

The software provided several geometric and topological manipulation algorithms for selected objects. Label combination algorithms merged multiple selected objects into single entities with automatic network table updates. Non-touching label splitting employed connected component analysis to separate spatially disconnected regions within labeled objects. Object deletion removed selected elements from both image data and network tables, with automatic topology updates.

#### Network Topology Modification

Network connectivity could be manually modified through link creation and deletion algorithms. Link nodes function created new network connections between selected nodes, while split nodes function removed existing connections. These operations provided manual network topology control with automatic updates to the network property tables.

#### Channel Override and Data Transfer

This method extracted highlighted regions from source channels and superimposed them onto target channels with automatic label transposition to prevent overlap conflicts. Empty channel handling created absorption arrays for isolated region extraction.

#### Spatial Measurement Tools

Integrated measurement algorithms provided distance and angular measurements with automatic scaling conversion. Distance measurements employed two-point linear algorithms with pixel-to-real-world conversion using user-defined scaling factors. Angular measurements utilized three-point algorithms with automatic acute angle calculation, where the second point served as the vertex. Measurement data was automatically tabulated with both voxel and scaled units.

#### Network Subgraph Isolation

Subgraph extraction algorithms created isolated network representations containing selected objects and their immediate neighbors. The method generated complete subgraphs placed in dedicated selection tables, enabling focused analysis of network subregions.

## Network Table Widget Right-Click Functions

### Table Sorting and Search

The network table implemented bidirectional sorting algorithms (ascending/descending) using any column as reference criteria. Integrated search functionality employed pattern matching with iterative result navigation through multiple instances of search terms.

### Spatial Navigation from Network Data

Reverse lookup algorithms enabled navigation from network table entries to corresponding spatial locations in image data. Single object navigation highlighted and centered individual nodes or edges, while pair navigation simultaneously highlighted connected node pairs and associated edge objects.

### Multi-Format Data Export

The software supported export to five standard network analysis formats: CSV for spreadsheet applications, Excel (.xlsx) format, Gephi (.gexf) format, GraphML for various network analysis platforms, and Pajek (.net) format. Each export maintained complete network topology and associated metadata.

### Network Table Management

Table management included bidirectional swapping between main network and selection tables, enabling detailed analysis of network subsets.

## Tabulated Data Widget Right-Click Functions

### Data Table Operations

Standard table manipulation included sorting and search functionality consistent with network tables. Export capabilities supported CSV and Excel formats for external analysis integration.

### Threshold-Based Node Filtering

Interactive thresholding algorithms utilized tabulated data to filter node populations based on numerical criteria. The method required structured input with integer identifiers and numerical values, enabling dynamic node selection based on analytical results.

## Example Data Analysis Methods

### Slime Mold Image

The slime mold image was obtained from the study by Tero et al^15^. The image was cropped to remove the large body of the slime mold in the upper right corner of the image. NetTracer3D’s machine learning segmenter was utilized on the color image to segment out the large feeding structures as nodes, then the inter-connecting pseudopods as edges, followed by thresholding out of small-object noise. With both loaded into their respective channels, the ‘Calculate Connectivity Network’ method was called with default parametes, except ‘Node Search’ set to 30 and ‘Edge Search’ set to 3. Further analysis shown included generating the network overlay from the ‘Overlays’ submenu, and Louvain community clustering and analysis of ‘Shortest Paths’, available from NetTracer3D’s ‘Analyze’ menu.

### Human Kidney Cortex Images

The images of human kidney cortices were taken from our previous light sheet study^31^, specifically, a ∼1mm^3^ region of cortex from reference sample ‘SK3’ of a 40-year-old male patient for the 20x, and a larger region of both cortex and medulla from reference sample ‘SK1’ of a 26-year-old female for the 5x. Segmentations were obtained from the previous study, which had been specifically created using Cellpose2^82^ to segment glomeruli, collecting ducts, and DAPI nuclei, as well as labkit^16^ to segment nerves.

### 20x

Glomerular and Collecting Duct segmentations were combined in NetTracer3D via the ‘Merge Labeled Images into Nodes’ function, then loaded into the nodes channel. The nerve segmentation was loaded into the ‘Edges’ channel, then a connectivity network was obtained with the ‘Calculate Connectivity Network’ function with default parameters, except ‘Node Search’ set to 10, ‘Edge Search’ set to 8, node labeling disabled (since the nodes had labels already), and image scalings set correctly. Radii data was generated with the ‘Calculate Radii’ function while glomerulus-to-collecting duct heatmaps were generated with the ‘Average Nearest Neighbors’ function, with the ‘Heatmap’ and Overlay 2’ params enabled, the ‘Use Centroids’ param disabled, the ‘Root Identity’ set to the Glomeruli, and the ‘Neighbor Identities’ set to the collecting ducts. The network overlay was generated from the ‘Overlays’ submenu. For Ripley’s clustering analysis of DAPI nuclei, a single glomerulus was used to mask its own cells. The segmentation of just this glomerulus was loaded into the edges channel, while the segmentation of its cells was loaded into the nodes channel. The ‘Ripley’s Clustering Analysis’ function was employed between the DAPI nuclei, with the ‘Define Boundaries How?’ parameter set to ‘Boundaries of Edge Image Mask’, which in this case contained the glomerulus segmentation, which restricted search radii to stay within this boundary. For Ripley’s clustering analysis of glomeruli, the same Ripley’s function was utilized on the entirety of the segmented glomeruli, but with the ‘Border Correction’ parameter enabled, which created mirrored nodes over the image borders, instead of using masked boundaries.

### 5x

A connectivity network was generated between glomeruli and nerves with the ‘node_search’ parameter set to 10 microns, the ‘label_nodes’ parameter disabled (as we had already labeled them), the ‘auto-trunk’ parameter enabled to simplify the node trunk, and the ‘fast_search’ parameter enabled. Betweenness Centrality was obtained using NetworkX.

### Mouse Kidney Cortex Image

Labkit^16^ was utilized to segment nerves and renin lineage cells. Cellpose3^11^ was utilized to segment glomeruli. The renin lineage cells and glomeruli segmentations were combined in NetTracer3D via the ‘Merge Labeled Images into Nodes’ function, then loaded into the nodes channel. The nerve segmentation was loaded into the ‘Edges’ channel, then a connectivity network was obtained with the ‘Calculate Connectivity Network’ function with default parameters, except ‘Node Search’ set to 10, node labeling disabled (since the nodes had labels already), and image scaling set correctly. Network graph visualization was done through the ‘Show Network’ function, while network overlays were generated from the ‘Overlays’ submenu. ‘Betweeness Centrality’ analysis was performed from the ‘Network Statistics Histograms’ menu, with high-value regions isolated via node-thresholding from the data table widget. ‘Glomerular’ and ‘JGA’ innervation was quantified with the ‘Calculate Node < > Edge Interaction’ function with ‘Node Search’ set to 10. For the branch-focused analysis, the nerve segmentation was first traced with the filament tracer to clean it up. The resulting filaments were labeled with the ‘Label Branches’ function with all default settings except enabling ‘Auto-Attempt to Reunify Main Branches’. The resulting branched edges were used to create a new network (with the glomeruli and renin masks again serving as nodes), with default parameters except ‘Node Search’ set to 10, node labelnig disabled, and the ‘Use Pre-labeled Edges’ parameter enabled. The resulting connectivity network retained branched edges whos shortest paths were evaluated with the ‘Nearest Network Neighbors’ function.

### Mouse Bronchus Image

Both bronchi and nerves were segmented with Labkit^16^. The bronchi segmentation was downsampled x5 in all dimensions. A combination of manual segmentation correction and NetTracer3D’s hole-filling algorithm was utilized to add the inside of the bronchi to the segmentation, as the initial stain only labeled epithelial regions. Small-object noise was thresholded out of the image, followed by an ‘Opening’ algorithm of 3-voxels in NetTracer3D, available from the ‘Clean Segmentation’ menu, as a means of further reducing noise and smoothign the segmentation. This smoothed segmentation had its branches labeled and a ‘branch-adjacency’ network generated via the ‘Calculate Branch Adjacency Network’ function, with all default parameters. To create the nervous network, the labeled branches from the ‘Opened’ segmentation were used to re-label the pre-Opened segmentation via the ‘Neighbor Labels’ function, with the ‘Labeling Mode’ parameter set to ‘Label Continuous Domains that Border Labels’, allowing the more spatially-accurate Segmentation to inherit the cleaner labels of the ‘Opened’ segmentation. This new segmentation was loaded into the nodes channel and upsampled to match the size of the non-downsampled nerve segmentation, which was loaded into the edges channel. Next, connectivity network generation was achieved via the ‘Calculate Connectivity Network’ function using default parameters except with node labeling disabled and image scalings set correctly. Community partitioning for both networks was done via Louvain clustering, network graphs were created from the ‘Show Network’ menu, and network overlays were obtained from the ‘Overlays’ submenu. Colored renders of branches were obtained from the ‘Color Nodes’ and ‘Code Communities’ functions.

### Angiogram Image

Blood vessels were segmented using NetTracer3D’s machine learning segmenter. The central brain arteries were isolated for use in network generation, removing other arterial domains such as meningeal arteries. Branch labeling and network generation was accomplished from the ‘Calculate Branch Adjacency Network’ function, with all default parameters except ‘Skeletal Voxel Branch Length to Remove’ set to 8. Community partitioning was done via Louvain clustering, network graphs were created from the ‘Show Network’ menu, and network overlays were obtained from the ‘Overlays’ submenu. Colored renders of branches were obtained from the ‘Color Nodes’ and ‘Code Communities’ functions. Betweenness Centrality was calculated from the ‘Network Statistics Histograms’ menu, with high-value regions isolated via node-thresholding from the data table widget.

### Lymph Node Nervous Image

The image was downsampled and segmented using NetTracer3D’s machine learning segmenter. This segmentation was then cropped to remove several hundred slices from the front of the Z-stack, which contained cross-labeled blood vessels as they were not the focus of this analysis. Branchpoint network generation was achieved via the ‘Calculate Branchpoint Network’ function with default parameters, except ‘Amount to Expand Nodes’ was set to 3. Branch labeling for the render, as well as branch length measurements, were obtained via the ‘Label Branches’ function with default parameters, except again setting ‘Amount to Expand nodes’ to 3. The rendering was obtained from the ‘Color Nodes’ function. Degree distribution, radial distributions, and nerve radii were obtained from their respective functions in the Analysis menu, with default parameters.

### CODEX Lymph Node Image

Cells were segmented using a Cellpose model trained to detect both nuclei (DAPI) and cell boundaries (NaKATPase). For the rest of the channels in the image, manual thresholding was performed to assign each cell their set of identities, alongside auto-computing the mean channel intensity per cell. A proximity network was generated for the 20 nearest neighbors within 200 pixels per cell using centroid based KDTree search. The node identities were used to assign neighborhood clusters, with the n_clusters parameter set to 10, and any node with fewer than 10 neighbors excluded. For the cell-based intensity clustering, KMeans clustering was used to cluster the cells based on their channel intensity profile, with n_clusters set to 10. UMAPs and Violin Plots showing identities were restricted to a smaller set of the available identities for visualization purposes.

### Stimulated-Raman-Spectroscopy Kidney Image

The four metabolite channels were brightness-normalized to make deeper Z-regions have comparable brightness to more shallow ones via NetTracer3D’s ‘normalize brightness’ function. NetTracer3D’s ‘Generate Hexagonal Nodes’ function was then applied to create hexagonal-prism nodes of base, length, and width of 8 microns within the masked boundaries of the tissue foreground. These nodes were assigned identities via manual thresholding. A proximity network was generated for the 20 nearest neighbors using centroid based KDTree search. The node identities were used to assign neighborhood clusters, with the n_clusters parameter set to 5. For the node-profiling, KMeans clustering was used to cluster the cells based on their Z-scored channel intensity profile, with n_clusters set to 5.

### Tumor Microspheroid Image

Hoeschst cells were segmented within Cellpose. These cells were assigned identities via manual thresholding across the three other channels. Proximity Networks were generated within 20 microns morphological borders using distance transform (which prevented network crossover through adjacent search regions). Louvain clustering was employed to create spatial network communities (as opposed to K-Means neighborhoods). The 3D nearest-neighbor heatmap was generated from NetTracer3D’s ‘Average Nearest Neighbors’ function, appraising the 5 nearest neighbors per cell starting from fibroblasts and looking for tumor cells. The nearest-neighbor distance matrix was also generated from the same ‘Average Nearest Neighbors’ method.

## Author Contributions

**S.J. and L.M.** conceived the idea; **S.J.** procured funding. **L.M.** wrote the first draft, generated the figures and performed all the example analysis; **L.M.** wrote the code and designed the program; **M.C.** and **S.S.** processed images; **P.R.K.** provided test CODEX datasets, oversaw collection of the tonsil image, and guided the direction of some of the analytical tools within NetTracer3D; **R.C.L.** imaged the tonsil; **R.J.S.** generated mouse lung LSFM data; **M.Y., L.M.L. and A.R.G**. generated mouse kidney LSFM data; **J.V.** and **L.S.** provided SRS datasets. All authors edited the manuscript. **S.J., L.M.** finished the final draft.

## Acknowledgements

We thank Xin Sun from UCSD for providing resources to generate the lung LSFM data. We are grateful to the Human BioMolecular Atlas Program (HuBMAP) for supporting this project and providing resources for data sharing. This work was in part supported by Human Biomolecular Atlas Program HuBMAP (U54DK134301 to S.J.) and Pediatric Center of Excellence in Nephrology at Washington University (P50DK133943 to S.J.). The content is solely the responsibility of the authors and does not necessarily represent the official views of the National Institutes of Health.

We thank Biospecimen and Laboratory Services at the University of Minnesota, including Andrew Nelson, Dina El-Rayes and Kira Krug for the Tonsil sample selection and procurement.

This work was supported by grants to Sanjay Jain from the NIH-funded Human Biomolecular Atlas Program HuBMAP (U54DK134301) and Pediatric Center of Excellence in Nephrology at Washington University (P50DK133943). The content is solely the responsibility of the authors and does not necessarily represent the official views of the National Institutes of Health.

## Data Availability

The image of the slime mold was used with permission from its publisher ‘The American Association for the Advancement of Science’, and can be found from the study by Tero et al^15^. The images of the human kidney cortex and 3D lymph node were obtained from publicly available datasets in the HuBMAP portal, available at [https://portal.hubmapconsortium.org/browse/publication/64ca708afbd3310816f8c7cedb389d5c] and [https://portal.hubmapconsortium.org/browse/dataset/b6eba6afe660a8a85c2648e368b0cf9f], respectively. The brain angiogram was obtained from a publicly available dataset from the study by Di Noto et al^83^, and can be found at [https://openneuro.org/datasets/ds003949/versions/1.0.1/file-display/sub-036:ses-20101030:anat:sub-036_ses-20101030_angio.nii.gz]. The mouse lung and kidney LSFM images are available at the following Zenodo repository (https://doi.org/10.5281/zenodo.17835509): [https://zenodo.org/records/17835509?token=eyJhbGciOiJIUzUxMiJ9.eyJpZCI6ImI5MmIyY2I0LWE1NTUtNGZmNy1iN2Y5LTY5ZWRiZTYxMTdkMyIsImRhdGEiOnt9LCJyYW5kb20iOiI5NGIyN2VhMTYwNDA2OWY4ZmE0YjFiMDk0ZjFiYTdjOSJ9.X6HKiy5Mexy2oAXfuGHdfFfqy_2Tknb1lDZPEqeEtd66gCEnAlZG6xGYR4ditD_NtQJPRAkbN3zuDseZF0TXYA]. The tonsil image can be found at this Zenodo directory: https://doi.org/10.5281/zenodo.18973389. The SRS image will be made available at a public HuBMAP portal upon publication in a peer-reviewed journal.

### Tutorial

For a tutorial on installing and using NetTracer3D, please reference the following Zenodo doi: https://doi.org/10.5281/zenodo.17873249

Make sure that the version of the Zenodo upload is set to its most recent to view the latest updated versions.

## Code Availability

NetTracer3D’s can be found most updated from its python package index page [https://pypi.org/project/nettracer3d/], where its source code is downloadable (commercial use may have fees – see below).

To install NetTracer3D as a python package, use the command ‘pip install nettracer3d’ in an environment with Python 3.11+ available. We recommend using anaconda to establish this environment, but it can also be done in a local environment on a computer. We recommend installing nettracer3d version 1.5.1+ for consistent behavior with the paper.

To install NetTracer3D to Windows directly with an installer, please download and run the latest release of the Windows installer at this github link: https://github.com/mclaughlinliam3/NetTracer3D/releases.

For more detailed instructions on installing and using NetTracer3D, please reference its documentation here: https://nettracer3d.readthedocs.io/en/latest/.

Commercial use is available for a fee. Copyright © is held by Washington University. Please direct all commercial requests for licensing, information, and limited evaluation copies to Washington University’s Office of Technology Management at.

## Information for Data Requests and Correspondence

Sanjay Jain

**ED Figure 1.**
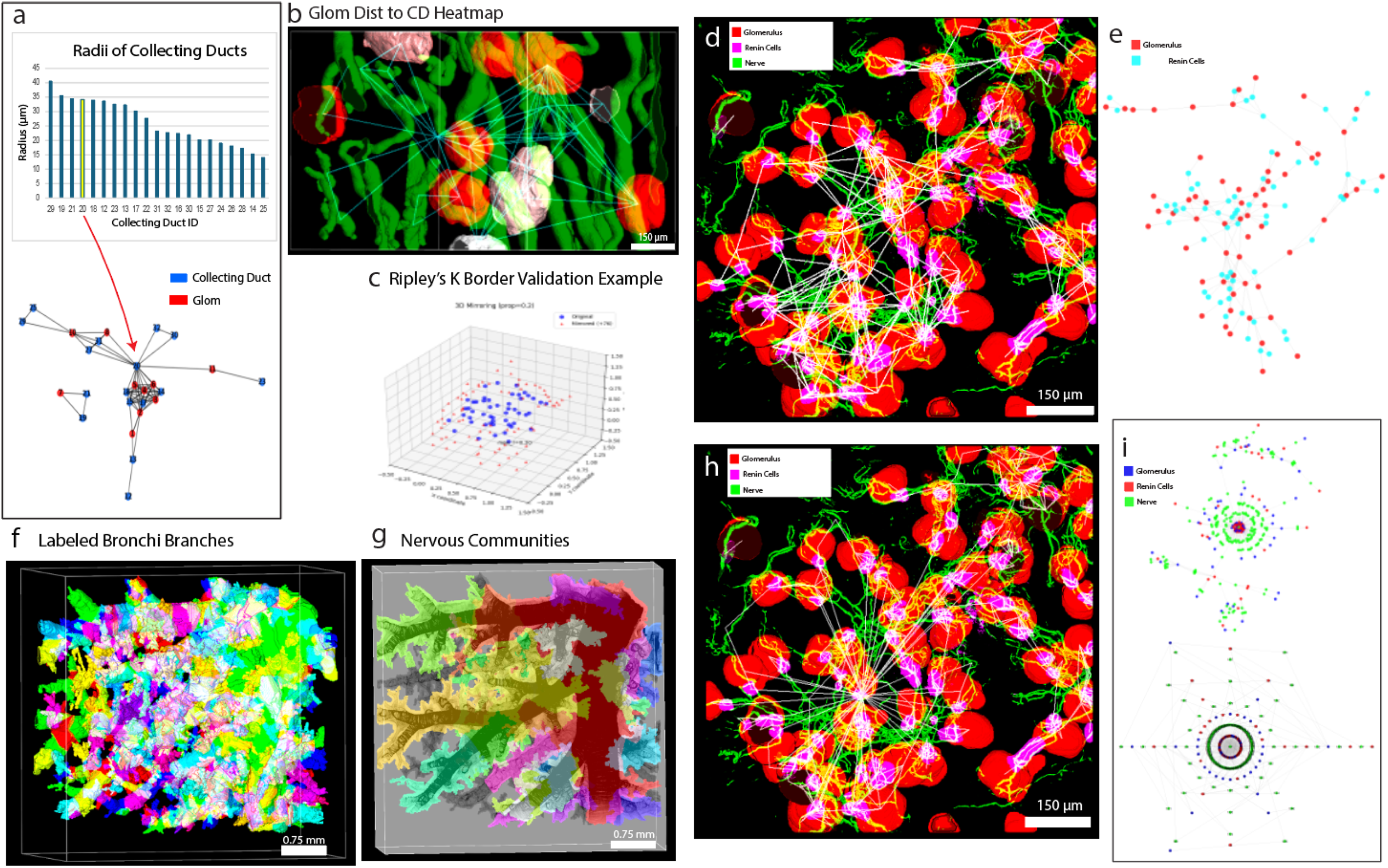
NetTracer3D Reveals Neural Connectivity Networks in Kidney and Bronchus of the Lung’. **a.** (Above): A bar graph showing the spread of the ‘largest radii’ of the collecting ducts shown in the human kidney cortex image from Figure 2a. (Bottom): A network graph of the glomerular-collecting duct neural network from the same image, as generated by NetTracer3D, with glomeruli shown in red and collecting ducts shown in blue. A centrally-located collecting duct (20) is indicated in the graph by the red arrow, as well as the yellow-colored bar in the radii graph, demonstrating the fourth-thickest collecting duct in the image. This particular duct demonstrates multiple ducts joining together, highlighted in yellow on Figure 2a. **b.** A heatmap with glomeruli encoded based on their distance to any collecting duct, demonstrating how NetTracer3D can be used for nuanced analysis of 3D relationships between structures. In this case, the red objects are quite closer to a duct while the white objects are an approximate theoretical average distance away. Greater-than-theoretical distant objects would be blue, although none were found in this instance. **c.** A graphical representation of how NetTracer3D enables border-safe Ripley’s Clustering Analysis. A set of blue theoretical nodes are arranged in 3D space. Around the borders of the image, a mirror set of red nodes have been created, allowing Ripley’s Clustering search radii to safely extend beyond the image border without showing falsely lowered clustering results. **d.** 3D render of the glomeruli/renin cells from Figure 2, now showing a connectivity network generated while enabling the ‘Auto-Simplify Trunk’ parameter which reduces the complexity of trunk elements. **e.** The network graph visualization from NetTracer3D for the network in Supplemental Figure 1d. **f.** The schematic of how the bronchi branches were labeled before being used to generate the neural connectivity network shown in Figure 2i. Each branch is indicated by a unique color. **g.** A 3D representation of the ‘Louvain Network Communities’ obtained from the bronchi-neural network. Different communities are shown by a unique color, overlayed on the original image of the bronchi (gray). Community domains roughly overlap with entire major branches, demonstrating a neural network that closely mirrors branch arrangement. Several branches that are not colored were not innervated by any nerves. **h.** 3D render of the glomeruli/renin cells from Figure 2, now showing a connectivity network generated while enabling the ‘Convert Edges to Nodes’ parameter which causes all edge plexuses to become nodes. **i.** Network graph visualizations from NetTracer3D for the network in Supplemental Figure 1h. (Top), Visualization using the spring layout. (Bottom), Visualization using the shell layout.

**ED Figure 2.**
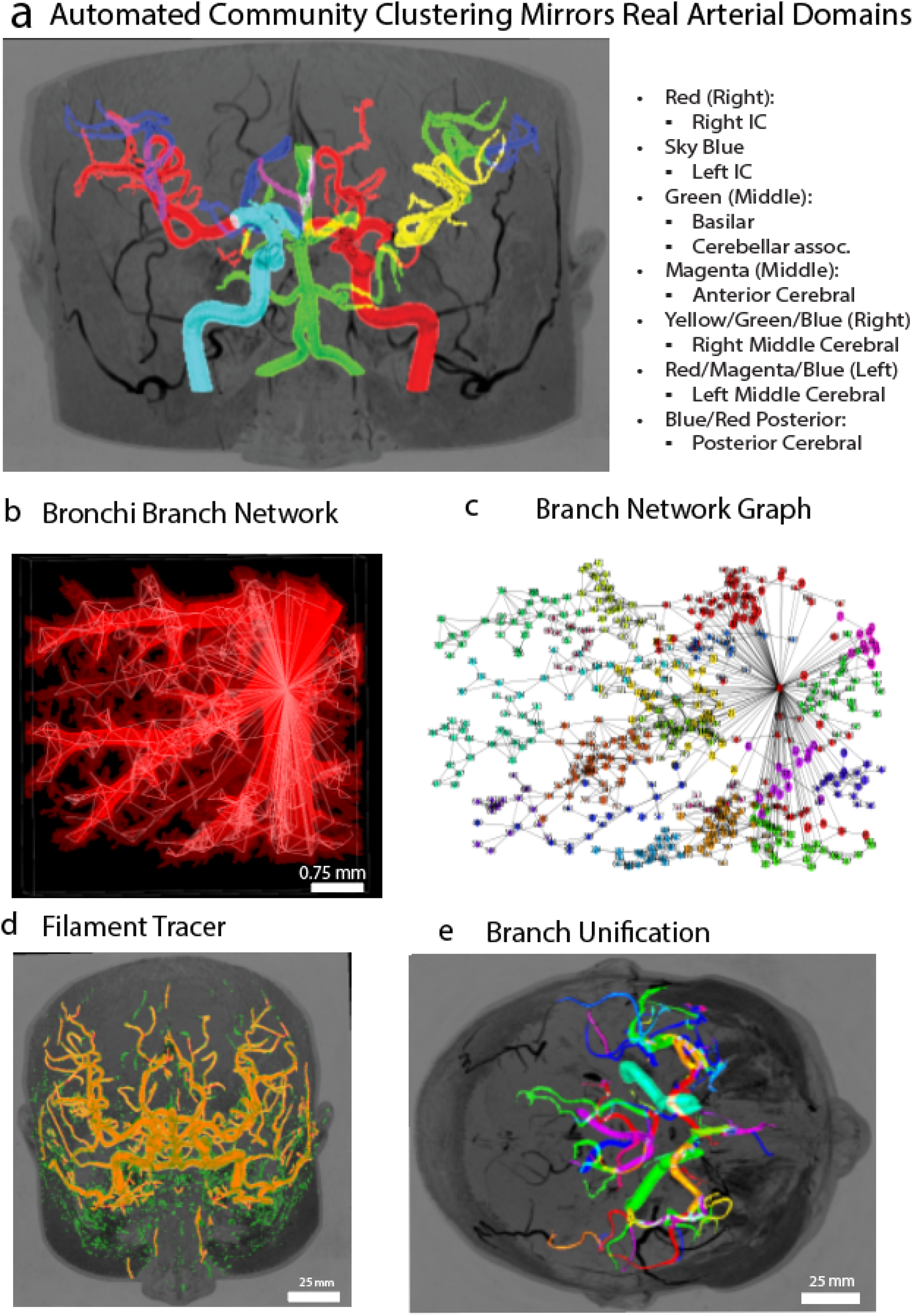
Branch networks of central arteries in the human brain and of neural patterns in a human lymph node’: **a.** The network communities of cranial vessels, from Figure 3b, now overlayed onto the original angiogram. The unsupervised Louvain clustering roughly mirrors known anatomical domains, demonstrating how unsupervised community detection often yields meaningful clusters. **b.** The bronchi image, previously used in Figure 2i to characterize neural patterns between branches, now used to make a direct network based on bronchial branch adjacency. Network connections between adjacent branches are shown with white lines. **c.** The network graph of the branches from Supplemental Fig 2b. Nodes are colored by community obtained from Louvain clustering. **d.** Example of the filament tracer being applied. The filament tracer attempts to trace filaments over segmentations of vessel or nerved based images, as a way to automatically clean up noise and smooth out surfaces. The angiogram from **Fig 3a** is shown alongside a messy segmentation (green), Overlayed is the result of the filament tracer on its default settings (red), demonstrating the removal of much of the noise and the bridging of gaps in the segmentation. **e.** An example of the branch reunification setting being applied. This optional setting will have branches attempt to reunify across branchpoint gaps if they can find branches of similar trajectories and radii, yielding a more natural branch-labeling schema where larger trunks are less likely to be subdivided into segments by branches coming off of them. In this instance, we apply this setting onto the segmentation from **Fig 3a**. Note how branches such as the trunk of the middle cerebral artery are no longer subdivided compared to **Fig 3a**.

